# A blueprint for biomolecular condensation driven by bacterial microcompartment encapsulation peptides

**DOI:** 10.1101/2024.11.13.623409

**Authors:** Daniel S. Trettel, Cesar A. López, Eliana Rodriguez, Babetta L. Marrone, Cesar Raul Gonzalez-Esquer

## Abstract

Bacterial microcompartments (BMC) are protein organelles with diverse metabolic capabilities. Their functional diversity is determined by an enzymatic core that is sequestered within a structurally conserved protein shell architecture. Segregation of protein cargo into the BMC is enabled by encapsulation peptides (EPs), which are short helical domains fused to core proteins through a disordered linker. Here, we investigate how EPs drive multicomponent cargo assembly into biomolecular condensates. *In vitro* experiments supported by molecular dynamics simulations demonstrate the importance of both conserved hydrophobic packing and electrostatic interactions in stabilizing trimeric EP bundles. Topological rearrangements of EP domains can drive programmable liquid-or gel-like partitioning *in vitro* and *in vivo*. This partitioning is found to be EP-specific, modular and can co-assemble at least three fluorescent reporters. In summary, we describe the molecular features necessary to drive biomolecular condensation using a widespread peptide tag. This work can serve as a blueprint for implementing EP biotechnology across diverse applications.

## Introduction

Bacteria are increasingly appreciated to dynamically organize their internal components.^1,2^ One important strategy for bacterial organization is the segregation of their metabolism into “organelles”, which can be found as lipid or protein membrane-bounded or unbounded (i.e. condensates).^3^ A widely studied prokaryotic organelle, called bacterial microcompartment (BMC), consists of protein polyhedra made of an outer protein shell which encases enzymatic cargo within.^4^ These organelles are widespread across taxa^5^ and can serve catabolic (called metabolosomes) or anabolic (called carboxysome) roles. BMCs have been proposed as a promising resource of structural (scaffolds, nanoreactors) and functional (substrate channeling, toxic chemical sequestration) motifs for their use in biotechnology.^6-10^ Recent work has connected that the biogenesis of some bacterial organelles go through phase-separated intermediates.^11-14^ This link ties protein condensation phenomena (specifically through liquid-liquid phase separation mechanisms [LLPS])^14^ to numerous critical cell functions like carbon fixation, cell division and organelle positioning.^1,15-23^

The early stages of BMC biogenesis have been suggested to make use of LLPS mechanisms, for example, in carboxysomes.^24^ In β-carboxysomes, the scaffold CcmM first nucleates liquid-like droplet formation^12,17,25^ of itself, Rubisco, and CcmN (i.e. procarboxysome)^26^ prior to shell recruitment by an encapsulation peptide (EP) in the C-terminus of CcmN.^27,28^ EPs are terminal amphipathic helices, typically 18 residues in length, fused to cargo by a poorly conserved disordered linker.^29-31^ The hydrophobic residues along one face of EPs have been shown to be a common motif critical for successful microcompartment packaging.^32^ This arrangement, as a single binding domain, does not strictly adhere to a canonical “stickers and spacers” (interaction motifs and linkers) framework commonly observed in biomolecular condensation.^33^ Metabolosomes have been similarly tied to using LLPS to guide assembly *in vivo*^34^, presumably through their EPs, resulting in the aggregation of protein “cargo” to each other and to the interior of the microcompartment shell. Notably, α-carboxysomes have been found to likewise utilize liquid intermediates to facilitate assembly by local condensation of the disordered scaffolds CsoS2, Rubisco, and shell components to trigger an assembly cascade without an identifiable EP.^13,18^

EPs have been observed to form *in vivo* aggregates^35,36^ but have not been explicitly studied for their ability to trigger biomolecular condensation of fused cargo^31^ for biotechnological applications. In this work, we study the self-assembly and condensation propensity of EPs. First, we demonstrate that the model EP from the aldehyde dehydrogenase (PduP) from *Salmonella enterica* fused to an mNeonGreen cargo reporter can trigger phase separation of fused cargo *in vitro*. Additional protein components can be assembled into condensed droplets if they contain an EP. *In vitro* experiments supported by molecular dynamics simulations demonstrate the importance of both conserved hydrophobic packing and electrostatic interactions in stabilizing trimeric EP bundles. Topological rearrangements of EP domains can exert control over liquid-or gel-like partitioning *in vitro* and *in vivo*. This partitioning is found to be EP-specific, modular and can co-assembly at least three fluorescent reporters into liquid-like foci in live bacteria. In sum, we provide a blueprint of the molecular features necessary to drive biomolecular condensation using the widespread EP peptide tag. This work paints a drastically new view of EP domains and demonstrates a new condensation system for the assembly of disparate metabolic machinery in bacteria.

## Results

### The PduP EP can drive cargo accumulation into biomolecular condensates

Our investigation started with the model PduP EP from the *Salmonella enterica* propanediol utilization (Pdu) BMC. This EP has been established to be important in cargo packaging within BMC shells^30^ and known to exhibit an α-helical structure by NMR and circular dichroism.^31,37^ We fused this 18 residue EP, along with its native 18 residue linker (Figure 1A), to the N-terminus of mNeonGreen^38^ (EP-mNG) to act as a reporter system and purified alongside wild type mNG. Biomolecular condensation is commonly triggered by adding molecular crowding agents to samples.^39^ As such, we titrated polyethylene glycol (average MN 2000; PEG2k) into solutions of mNG and EP-mNG at 20 µM and measured sample turbidity as a proxy for aggregation. Turbidity is commonly used to assess biomolecular condensation as condensed particles refract light and the resulting signal serves as an indirect measure of their number and size.^40^ We observed no changes in turbidity for mNG while EP-mNG turbidity increased in a sigmoidal fashion with an inflection at 17.2% ± 0.6% (w/v) PEG2k (Figure 1B). We then chose 20% (w/v) PEG2k as a constant and assayed protein concentration for turbidity changes (Figure 1C). Again, mNG turbidity remained constant while EP-mNG increased with an approximate inflection at 33.1 µM ± 0.4 µM.

**Figure 1:**
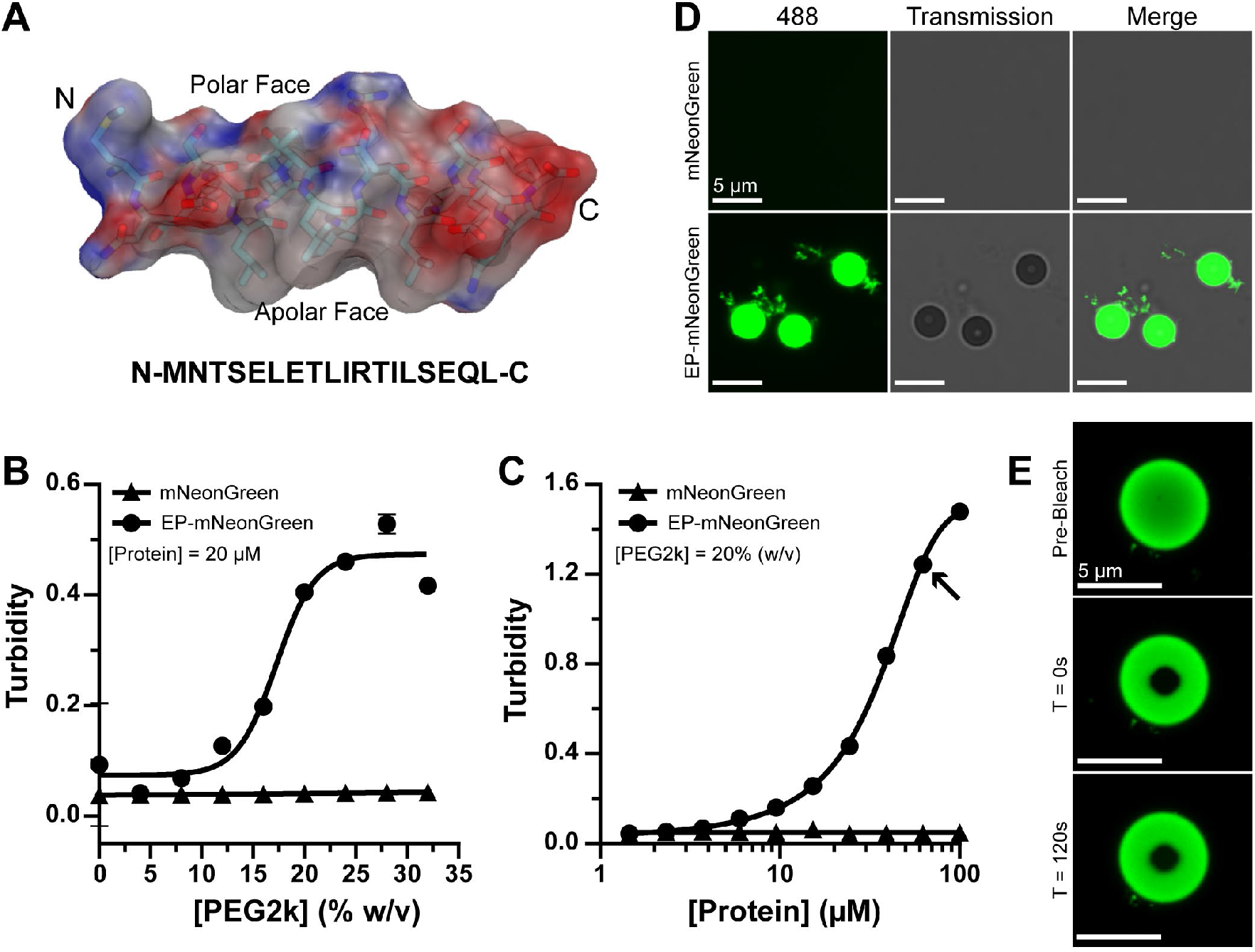
The encapsulation peptide domain from PduP triggers biomolecular condensation of cargo. **(A)** The model EP from PduP is an amphipathic, 18 residue α-helix fused to amino terminus of protein cargo by a disordered linker. The residues depicted in the helix representation are in black. Turbidity assays are used to assess particle formation from EP fusions as a function of **(B)** molecular crowder PEG2k and **(C)** protein concentration. The arrow in Panel C are samples taken for imaging. **(D)** Confocal imaging reveals the presence of spherical particles in solution for samples containing EP fused cargo. **(E)** Photobleaching experiments show minimal/no fluorescence recovery.

The above data suggested that the EP tag can result in particle formation in solution. However, turbidity assays cannot differentiate between cargo aggregation and condensation.^41^ We investigated this by imaging turbid samples (Figure 1C, arrow) with laser scanning confocal microscopy. Imaging showed no structures formed by mNG itself (Figure 1D). In contrast, EP-mNG was able to spontaneously demix and form spherical droplets in solution (alongside some amorphous material) which is suggestive of phase separation coupled to percolation (PSCP).^42^ Biomolecular condensates that form through PSCP should display changes in their viscoelastic material properties as a function of time.^42^ We tested the material state of these droplets by performing a fluorescent recovery after photobleaching (FRAP) experiment to monitor the recovery of fluorescence.^43^ If liquid-like, droplets will experience quick recovery of fluorescence (typically <60s timescale) in bleached zones owing to rapid exchange and equilibration of bleached and unbleached molecules. The droplets formed by EP-mNG, however, did not experience significant recovery over a qualitative 2-minute sampling (Figure 2E) indicating slow dynamic internal rearrangements indicative of a gel-like state. The droplets were further commonly observed in arrested fusion states that never resolved (Supplemental Figure 1A), and likewise reminiscent of gelation. We also found that if our usual order of component addition was inverted (protein from a concentration stock added last) then these mid-fusion assemblies or larger aggregates, which appear as numerous unresolved droplets, were more commonly observed (Supplemental Figure 1B). These data suggest that these assemblies undergo rapid networking which cannot sufficiently resolve into distinct droplets when triggered from a concentrated protein stock. This, and their inability to recover quickly from bleaching, suggests that EPs enable PSCP to form physical microgels and that transitions in their viscoelastic properties happen on relatively short timescales.

**Figure 2:**
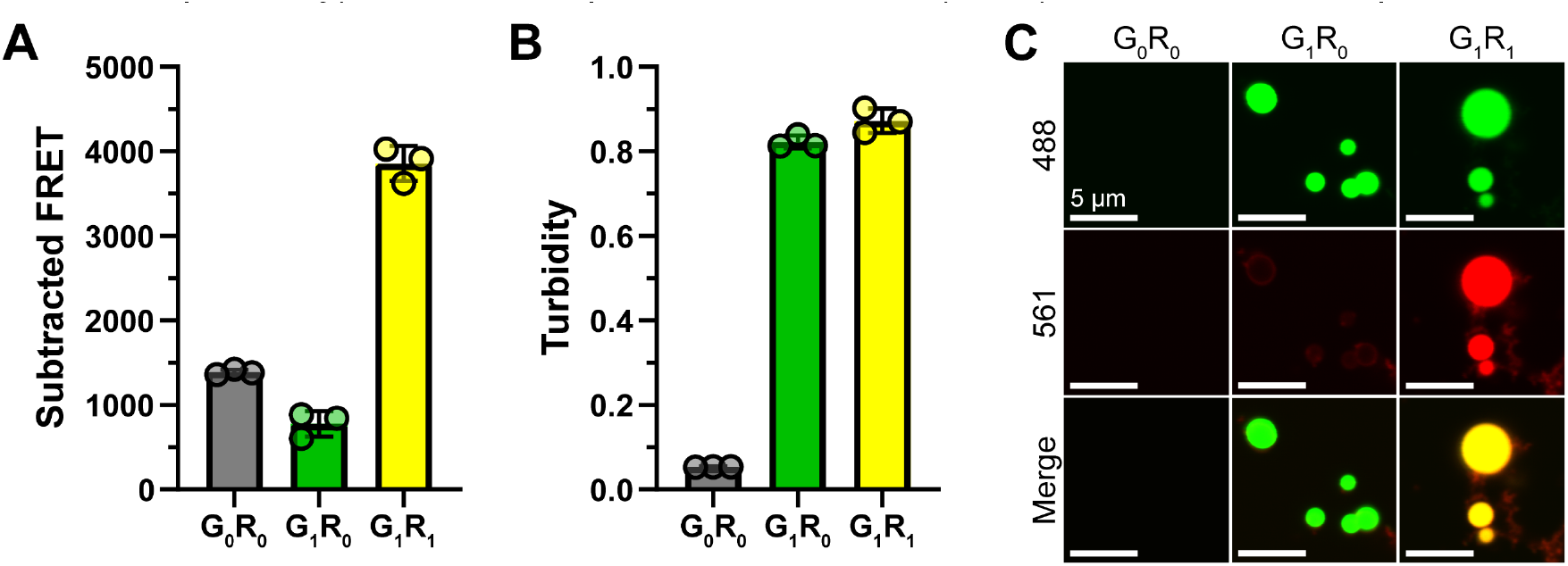
Specific *in vitro* co-assembly of multiple cargoes with encapsulation peptides. **(A)** Fluorescence resonance energy transfer (FRET) was used to assess assembly of a donor mNeonGreen (G) and acceptor mScarlet-I3 (R) with (1) or without (0) N-terminal EP fusions. FRET was calculated by subtracting the emission of mNeonGreen by itself at the peak acceptor emission wavelength (590 nm). Significant FRET only occurred when both cargoes have encapsulation peptides. **(B)** Turbidity assays of samples confirm particle formation only when at least one cargo had an EP. **(C)** Confocal imaging showed that specific co-assembly of mNeonGreen and mScarlet-I3 only occurred when they both had EP fused to their N-termini.

### The EP domain can drive specific co-assembly of multiple species

We next sought to understand if EP-driven particles can include more than one component. We tested this by mixing purified mNeonGreen (G) and mScarlet-I3^44^ (R) either with (G_1_ or R_1_) or without (G_0_ or R_0_) an EP fusion and measured the results by fluorescence resonance energy transfer (FRET), turbidity, and imaging methods (Figure 2A-C, respectively). Mixed samples with no EPs (G_0_R_0_) were unable to produce a significant FRET or turbidity signal and confocal imaging found no particles in solution. Samples which included an EP only on mNeonGreen (G_1_R_0_) showed a slight decrease in FRET signal and high turbidity. Imaging results indicated that the turbidity was a result of mNeonGreen particles with a weak outer halo of mScarlet-I3. The slightly lower FRET signal was likely due to mNeonGreen condensation in solution which limited mScarlet-I3 interactions. Lastly, samples with EPs on both proteins (G_1_R_1_) produced the highest FRET signal and turbidity equal to G_1_R_0_ samples. Confocal imaging showed that G_1_R_1_ particles were formed from co-assembled mNeonGreen and mScarlet-I3, which agrees with FRET data. These data demonstrated that EPs drive specific co-assembly of multiple cargoes *in vitro*.

### EPs assemble through hydrophobic packing and a specific salt bridge

Our model system gives us an opportunity to screen EP self-assembly through simple turbidity measurements. EPs have traditionally been thought to act solely through relatively non-specific hydrophobic packing interactions predominantly along their apolar face (Figure 1A).^32,45^ We confirmed this by performing molecular dynamics (MD) simulations by inspecting the dimerization process of two EP helices. A residue contact map of tightly associated helices agrees with the notion that hydrophobic residues, for instance I10, form the closest and most abundant contacts between bound helices (Figure 3A, black boxes) and have been previously implicated with being critical for cargo encapsulation within microcompartment shells but not self-assembly.^32^ A closer look at simulated EPs also identified an antiparallel salt bridge network formed by E7 of one helix and R11 from the other spaced by exactly one helical repeat (Figure 3B). We tested the importance of these residues by making several point mutations and compared their condensation propensity as a function of NaCl concentration (to disturb the salt bridge) and protein concentration. The mutations we chose to assay are R11K (conserved mutation), R11A (removal of salt bridge), E7D (side chain one carbon shorter), and I10S (disrupt hydrophobic core).

**Figure 3:**
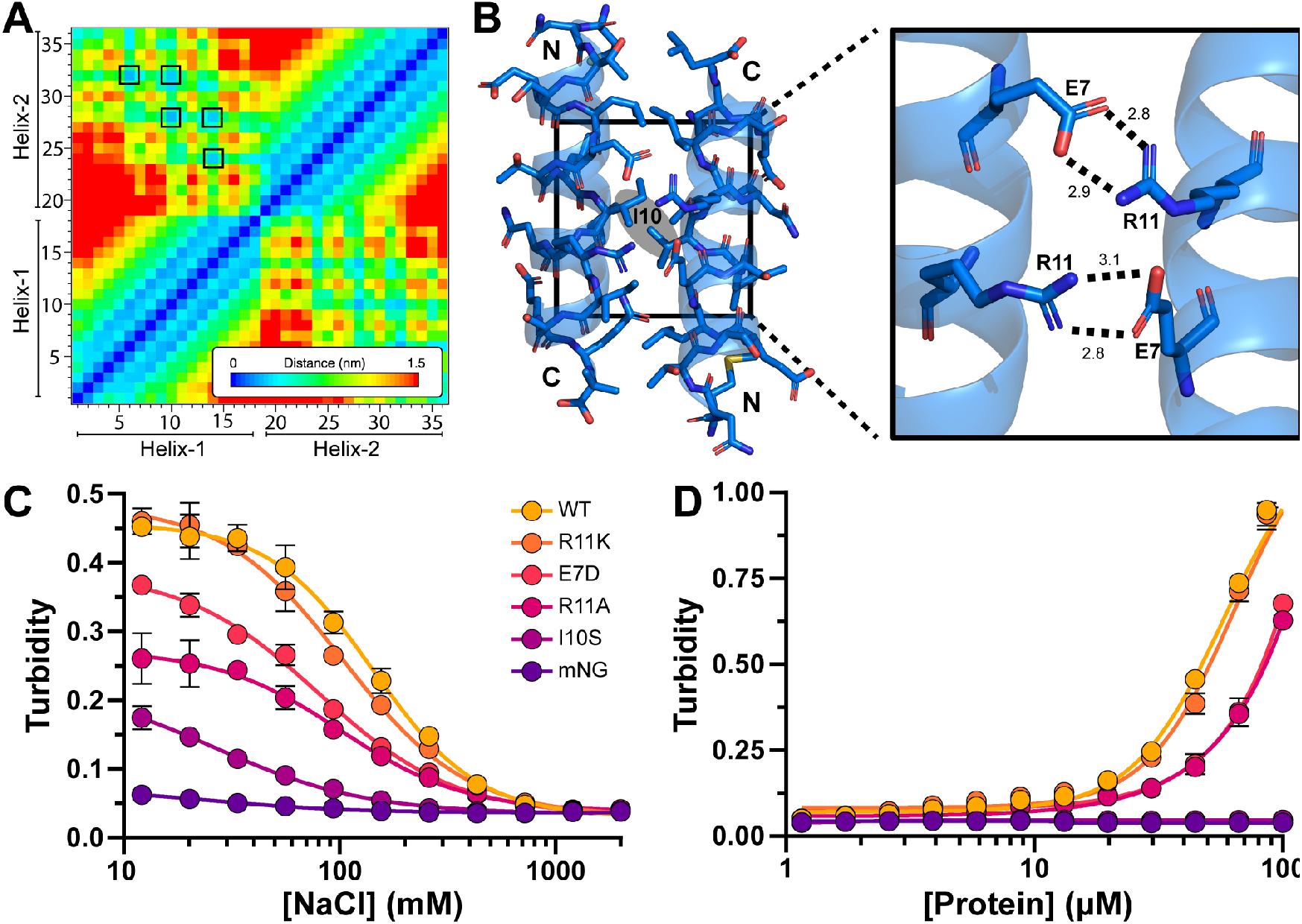
Encapsulation peptides rely both on hydrophobic packing and a newly identified salt bridge network. **(A)** A distance matrix of two assembled PduP EPs shows that the closest interactions are from hydrophobic residues (black outlines) along the apolar face. **(B)** Computational models of EP dimers suggest that they prefer an antiparallel orientation which results in a salt bridge network between E7 and R11. Several mutants were made, and their condensation propensities were screen by measuring turbidity as a function of **(C)** NaCl for electrostatic screening and **(D)** total protein concentration. These data demonstrate that EP self-assembly is sensitive to ions and condensation can be negatively impacted by disrupting the salt bridge and hydrophobic core.

A salt titration assay confirmed that all EP mutants, to varying extents, resulted in turbidity and that turbidity is negatively impacted by increasing salt (Figure 3C). At low salt saturation, the R11K mutant performed similarly to the wild type PduP EP confirming that functionality is conserved. However, as expected by its hydrogen bond strength, high salt concentration had slightly more effect on the R11K mutant. The E7D mutant, which may simply weaken the salt bridge, was less robust than the R11K mutant but more robust than the R11A mutant which completely removed that interaction. Further, the inflection points of all salt bridge mutants were shifted left towards lower concentration (140 mM for WT versus 80-100 mM for mutants), revealing a heightened salt sensitivity. These trends were further reflected when titrating protein concentrations (Figure 3D). Again, WT and R11K performed similarly while salt bridge mutants showed an intermediate self-assembly propensity. The I10S mutant had no assembly propensity. These results indicate that the computationally predicted salt bridge is real and impacts self-assembly strength of EP domains. The most critical residue we tested, however, is I10 within the hydrophobic core.

### Metabolosome EPs are functionally distinct from carboxysome EPs

Most BMCs use encapsulation peptides to accumulate cargo within their shells.^29^ This includes β-, but not α-, carboxysomes which use the C-terminus of the protein CcmN to drive associations between CcmM-RbcLS coacervates and the shell.^28^ The C-terminal extension of CcmN proteins have been regarded as EPs because they literally link cargo-shell interactions. This is a very different arrangement for cargo inclusion interactions compared to metabolosomes, wherein EPs typically exist fused to enzymatic cargo as opposed to CcmN which acts as a discrete intermediary between enzymatic cargo and the shell. Sequence comparisons of several candidate metabolosome and CcmN EPs showed that metabolosome EPs largely conserve the salt bridge related residues we have identified with a sequence spacing corresponding to one helical repeat (Figure 4A). The EPs of CcmN, however, do not carry these sequence elements. We investigated this further by screening the ability of different EPs fused to mNeonGreen to trigger biomolecular condensation of this cargo. In addition to our PduP model, we also selected PduD (swapped charges), PduE (reportedly not sufficient for cargo encapsulation^46^), EutC (different BMC), and CcmN (from *Synechococcus elongatus* PCC 7942).^26,28^ All metabolosome EPs (PduDEP, EutC) were found to result in cargo condensation to varying degrees (Figure 4B). PduD was the most robust followed by PduP, EutC, and finally PduE. Meanwhile, the CcmN EP was not able to trigger significant cargo aggregation. Protein droplets were likewise found to exist in all metabolosome EP samples, but not with CcmN (Figure 4C). These self-assembly trends were likewise recapitulated with *in vivo* overexpression, with CcmN unable to drive cargo partitioning (Supplemental Figure 2A, 2B).

**Figure 4:**
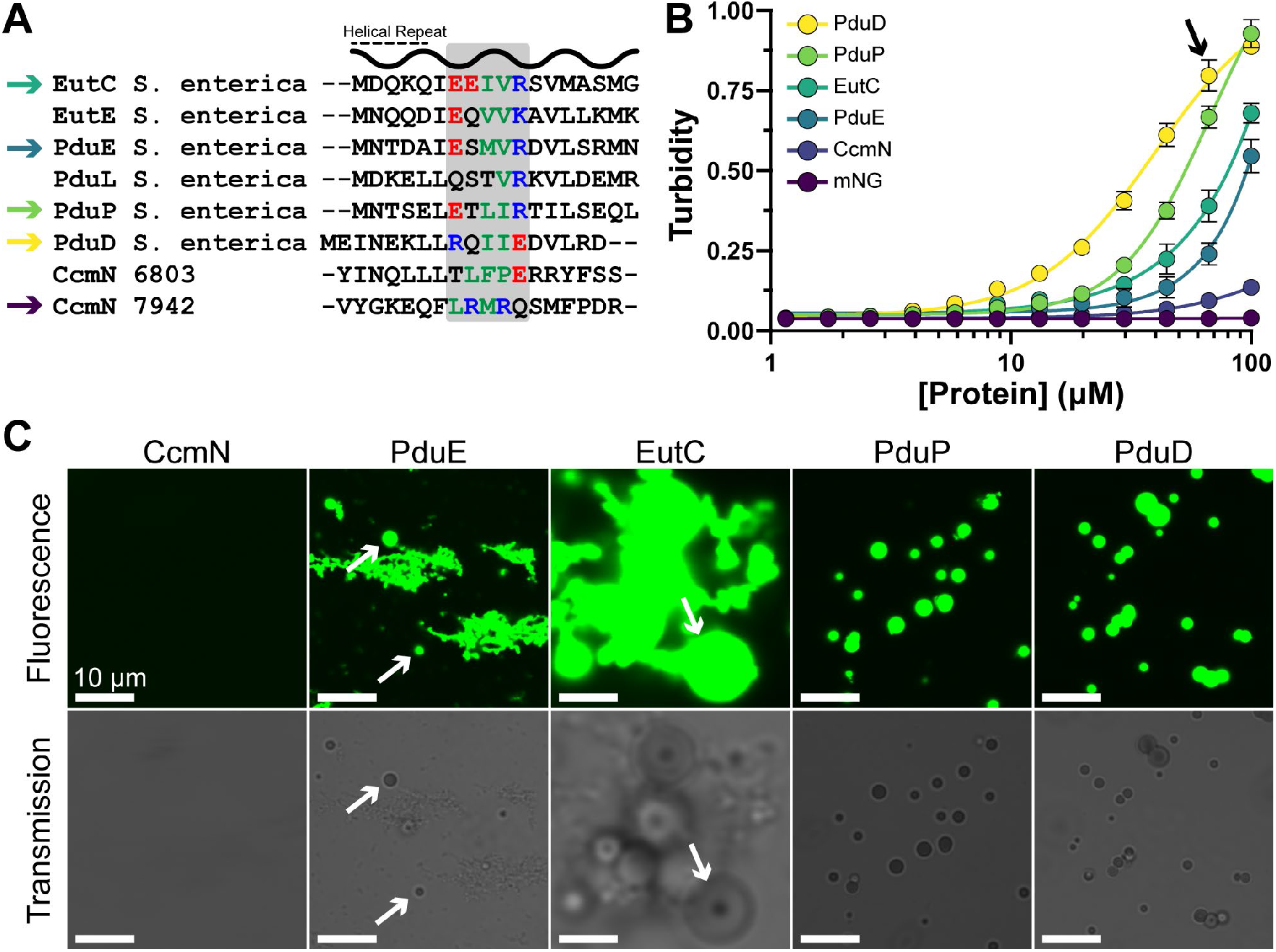
Metabolosome EPs have distinct sequence elements which carry functional consequences. **(A)** An alignment of several EPs revealed that metabolosome EPs carry oppositely charged residues separated by one helical repeat. These elements are not maintained in carboxysomal CcmN sequences. **(B)** A turbidity assay demonstrated that these sequence differences result in CcmN being unable to condense cargo under conditions that trigger condensation for metabolosome EPs. **(C)** Aliquots from turbid samples (arrow in panel B) were imaged with confocal microscopy. Spherical droplets were observed in all metabolosome samples, to varying extents (arrows) and most clearly defined in PduP and PduD samples. CcmN did not drive droplet structures.

### EPs associate as trimeric complexes

Our computational calculations demonstrated that EPs can form a specific salt bridge when paired in an antiparallel orientation (Figure 3B). This association, however, splays open the hydrophobic core between the helices, indicating additional association capacity. We experimentally tested this by fusing serial repeats of the PduP EP and its native linker (EP1, EP2, EP3) to the N-terminus of an mNeonGreen cargo (Supplemental Figure 3A). We hypothesized that this synthetic arrangement of intrinsic valency, mimicking a more classic stickers and spacers arrangement, will either (1) increase binding capacity linearly or (2) form intramolecular arrangements (due to proximity and connectivity) that self-quench binding capacity and lead to diminishing returns. We assayed these designs for self-assembly propensity and found that the EP2 design moderately outperformed the EP1 while the EP3 performed similarly or slightly worse than the EP2 (Figure 5A). Specifically, curve inflections for the EP1, EP2, and EP3 designs were found to be (in µM) 33.1 ± 0.4, 15.5 ± 1.0, and 19.7 ± 1.9 respectively. The morphology of all samples was also similar, and all shared the ability to form droplets across a wide range of concentrations (Figure 5B). Here, increasing protein concentration increased both the size and quantity of droplets from all designs. These designs were similarly assayed for response to PEG2k concentration with identical trends observed (Supplemental Figure 3B). These results demonstrated intramolecular valencies above two quenches additional associations, thereby providing evidence for associations greater than a dimer.

**Figure 5:**
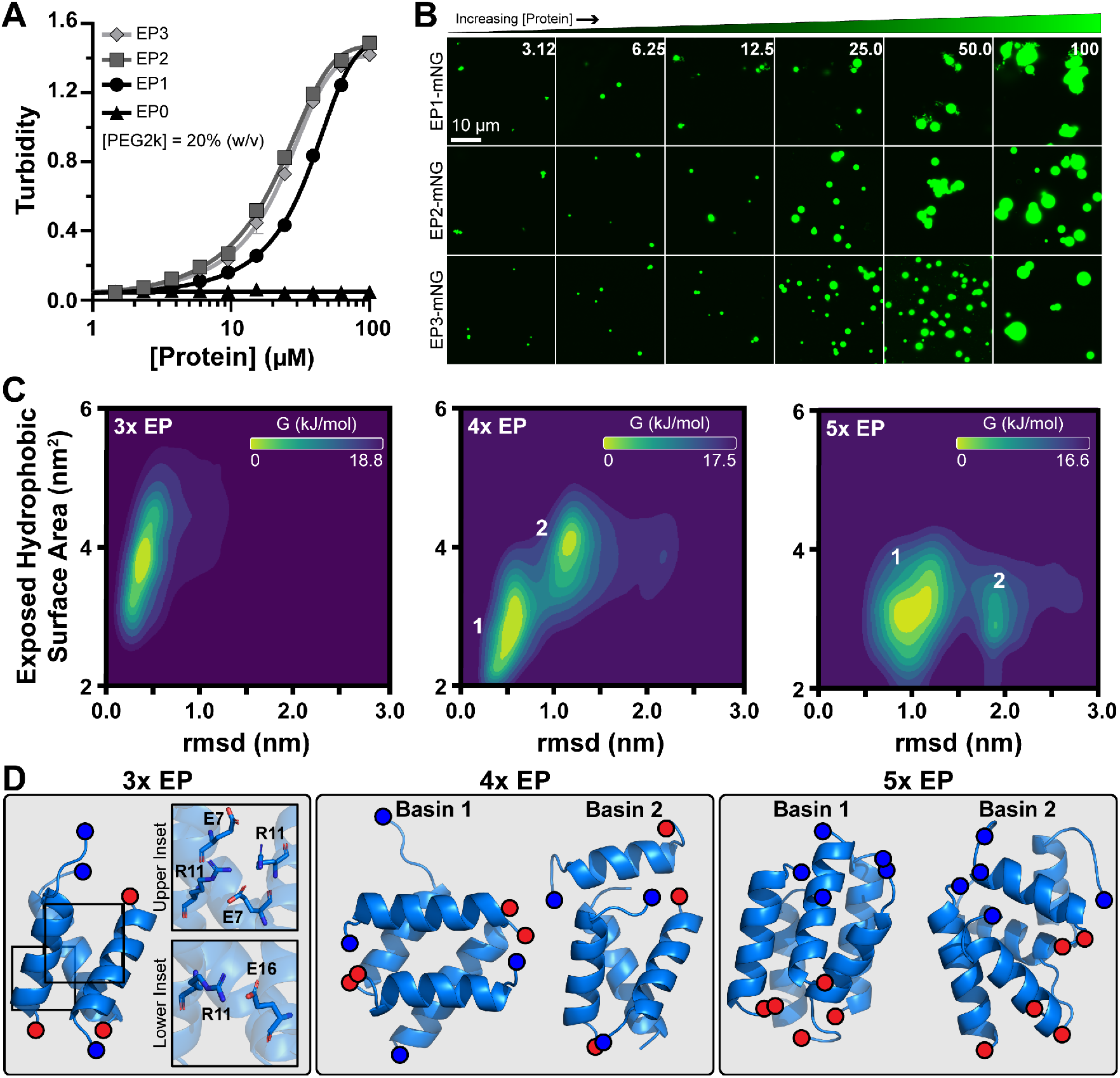
Encapsulation peptides self-assemble in a multivalent manner. **(A)** Turbidity was measured as a function of protein concentration for synthetic EP designs where EPs were fused in serial to the cargo N-terminus. Additive copies had diminishing results. **(B)** Despite differences in sequence topology and condensation propensity, confocal imaging showed all designs formed similar particles in solution across a range of concentrations (in µM). **(C)** Computational simulations were conducted with three, four, or five EP helices to study their preferred assembly state as a function of exposed hydrophobic surface area and root mean square deviation (rmsd). Simulations with three EPs coalesced to one basin with low rmsd while those with four or five showed two dominant populations. **(D)** Structural analysis of these basins revealed that EP assembly is associated salt bridge interactions along their solvent exposed surfaces.

To understand the higher order association mechanism, we resorted to MD calculations to analyze the saturation binding capacity of EPs. We ran simulations of 3, 4, or 5 associated PduP helices and summarized their dynamics and energetics as a function of both positional variance (RMSD) and exposed hydrophobic surface area (Figure 5C). Simulations with 3 helices coalesced towards one single basin while those with 4 or 5 helices formed two. Representative configurations for the different basins are provided in Figure 5D, highlighting the helical association. In particular, the 3-helix configuration remained very stable and formed a tightly packed bundle. In addition to the hydrophobic interaction, a close inspection indicates that this structure is further stabilized by salt bridges formed by pairs E7-R11 and R11-E16 (Figure 5D left panel). In contrast, other stoichiometries (4 and 5 helices) led to unstable bundles that transition between configurations. In both cases a trimeric bundle was observed in basin 2 which led to a single or dimeric helix that is unstably associated. These insights led us to conclude that optimal trimeric bundles are expected to guide the association of EP domains. These results may explain the condensation results of our tandem sequence designs. For instance, the EP2 construct would always have additional networking capacity even if the domains intramolecularly dimerize. The EP3 design, in contrast, may be more likely to intramolecularly trimerize *in vitro* and block further network extension. These results confirmed that EP association is not only dependent on hydrophobic complementarity, but clearly by specific, highly conserved interactions that lead to a trimeric assembly preference as opposed to general aggregation.

### EP topology can alter partitioning and material state in vivo

Our *in vitro* model system thus far has included strictly monomeric proteins, which differs from those natively tagged with EPs and may not reflect physiological reality. For instance, PduP is a homotetramer^47^ and PduCDE, which has EPs on PduD and PduE, is a dimer of trimers.^48^ We investigated the effects of binding site valency by again implementing our serial EP designs fused to a monomeric mNeonGreen against a tetrameric E2-Crimson^49^ (hereafter Crimson) to differentiate between intrinsic and emergent valency, respectively (Figure 6A). We sought to explore the functional differences between these two arrangements *in vivo* to gain a proper understanding of EP-driven assembly in living cells. Accordingly, we overexpressed these designs in *E. coli* and used laser scanning confocal microscopy to image their phenotypes *in vivo* (Figure 6B). Both the wild type mNeonGreen and Crimson showed a uniform fluorescence inside of cells. Meanwhile, increasing the EP copy number on mNeonGreen similarly increased the segregation of multiple (typically 2+) mNeonGreen cargo foci within the cells. This effect was also observed with a single EP copy fused to Crimson cargo, which formed single elongated foci. We quantified this effect by calculating a partition ratio (the intensity of background subtracted foci divided by non-foci areas of the same cell) which is analogous to how much denser foci are compared to the dilute areas of the cell (Figure 6C). Quantification in this way gives the mNeonGreen and Crimson controls a median partition ratio close to 1.0, indicating uniform fluorescence distribution throughout the cell. This analysis further shows that, in contrast to our *in vitro* tests, additive copies of EP domains increased partitioning up to 9-fold over the mNeonGreen control for the EP3 design. This effect was exacerbated in the EP-Crimson design which were able to achieve a median partition ratio of over 500, making it nearly all-or-nothing. Overall, these results demonstrate the importance of valency in driving cargo networking and that our serial repeat design likely prefers intermolecular, and not intramolecular, associations *in vivo*. We believe these differences compared to our *in vitro* experiments may stem from the dense cytoplasmic milieu inhibiting specific intramolecular associations and promoting intermolecular networking.

**Figure 6:**
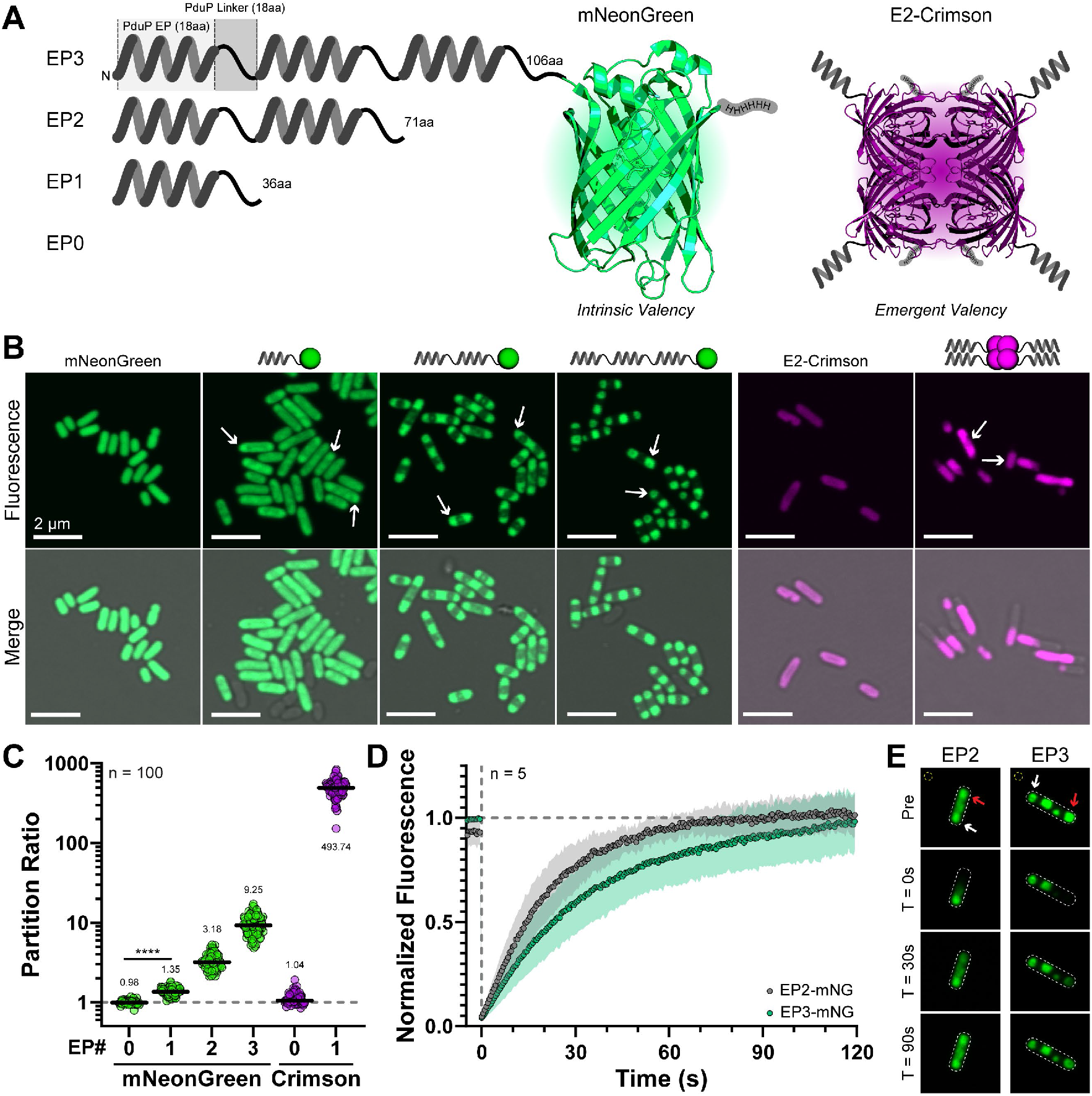
Valency effects cargo partitioning in vivo. (A) We tested the cargo partitioning performance of serial EP designs on a monomeric mNeonGreen cargo (intrinsic valency) and on a tetrameric E2-Crimson cargo (emergent valency). The mNeonGreen structure is from PDB 5LTR and E2-Crimson is represented by a tetrameric DsRed 1GGX. **(B)** Overexpression of designs for intrinsic valency showed that increasing valency increased partitioning ability. The emergently valent EP1-Crimson showed elongated single foci. Arrows denote several example foci within each sample. **(C)** Partitioning was calculated by the quotient of background subtracted foci signal over non-foci signal. This analysis confirmed that partitioning increases with additional EP domains and that emergent valency outcompetes intrinsic in all cases. Medians are denoted within each sample. **(D)** Fluorescence recovery after photobleaching (FRAP) experiments showed that the EP2 and EP3 designs differed in their recovery rates, but both were liquid-like. **(E)** FRAP was calculated by normalizing the background subtracted (yellow circle) intensity of bleached foci (red arrow) to another unbleached foci (white arrow) within the same cell

We next assayed the material state of EP-driven foci within cells by measuring *in vivo* FRAP.^43^ Here, single foci within cells were bleached with a laser pulse and their fluorescence recovery was monitored with 0.5 second intervals. FRAP experiments were focused on comparing the relative differences between EP2 and EP3 designs (Figure 6D) due to quick recovery of EP0 samples and EP1 foci being hard to find. FRAP was calculated by normalizing the background (yellow circle, Figure 6E) subtracted fluorescence of bleaching foci (red arrow) to an unbleached focus (white arrow) to see when separate foci equilibrated. The EP2 and EP3 designs were able to achieve full re-equilibration relative to unbleached foci within 2 minutes and with half-times of 16.2 s ± 8.5 s and 27.2 s ± 14.6 s respectively. We note that this is slower than recovery of free fluorescent protein, which is on the order of a few seconds.^50^ This contrasted drastically with results from cells expressing EP-Crimson, which demonstrated a linear and incomplete recovery profile within this timeframe, indicative of a gel-like state (Supplemental Figure 4).^22,23,43^ The relatively large errors may stem from different distances between bleached and unbleached foci within cells, therefore leading to longer diffusion distances. Despite this, the recovery curves of EP2- and EP3-mNG were significantly different (p < 0.001). This experiment alone cannot determine if the difference between the EP2 and EP3 designs stem from diffusion rate differences (EP3 is larger and may migrate slower) or due to slight changes in their condensed material states. However, it would make sense that the EP3 does equilibrate slower than EP2. The EP3 design likely forms more intermolecular contacts to shift equilibria, a function of on and off rates, as informed by its increased partition ratio and thereby slowing equilibration. These results demonstrated that EP-driven condensation in bacteria was liquid-like in contrast to our *in vitro* results; and in equilibrium with the excluded cytosol, and not inclusion bodies, which have been demonstrated to not recover during similar experimentation.^43^ Further, topological rearrangements of EPs can be used to encode for different cytosolic behavior.

### EP partitioning is specific and can partition multiple components in vivo

Overexpression of a single protein fused to the PduP EP can partition it between a condensed and dilute phase *in vivo*. However, more complex arrangements exist naturally, and multicomponent systems would be desired synthetically. We first tested the ability of a single mNeonGreen cargo to partition when co-expressed equally (assumed from ribosomal binding sites) with mScarlet-I3 and mTagBFP2.^51^ Designs were made with mNeonGreen encoding for up to three EP repeats and given a nomenclature based on the gene ordering and number of EP repeats (Figure 7A, top). For instance, a design with two repeats only on mNeonGreen is named G_2_R_0_B_0_ (for green with two EPs, red with zero, blue with zero). Constructs were overexpressed and mNeonGreen partitioning was calculated as before. The general trend of more EP repeats leading to greater partitioning was again observed (Figure 7B, 7C). However, the effect was muted compared to the single-expression designs which we suspect stemmed from a competition for resources between the three proteins during expression. That is, the competition may have lowered the effective expression, and therefore concentration, of EP domains and lessened mNeonGreen was observed again confirming the specificity. In addition, the G_3_R_0_B_0_ expressing cells also reveal dark spots in the red and blue scans where the mNeonGreen appeared to localize, suggesting exclusion of non-EP tagged proteins.

**Figure 7:**
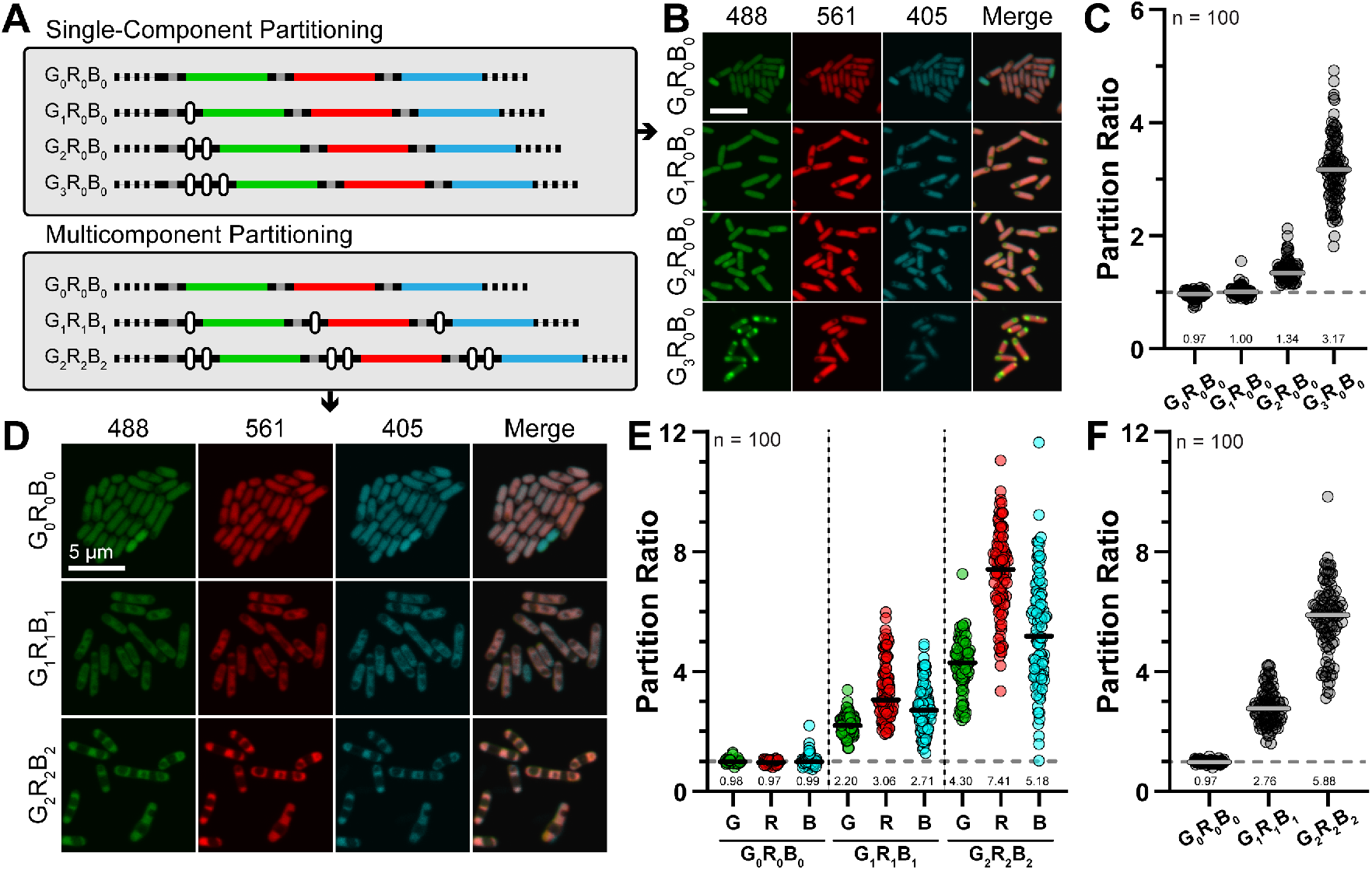
Programmable and specific multicomponent cargo partitioning using the encapsulation peptide domain. **(A)** Synthetic operons were designed with three different fluorescent cargos of mNeonGreen, mScarlet-I3, and mTagBFP2 to test specific single and multicomponent partitioning. These designs were given a nomenclature based on the gene order and number of EP domains fused to that cargo. Here, EP repeat elements are signified by white rounded rectangles and ribosomal binding sites by gray lines. **(B)** Overexpression of designs expressing EP domains only fused to mNeonGreen showed only green components in foci formed. **(C)** As in the single-protein designs, more EP domains resulted in increased partitioning of the EP cargo. **(D)** Fusing EP domains to all cargo resulted in their co-assembly into condensed “zebra-striped” bands within the cell. Calculating the partitioning of these cargoes either **(E)** separately or **(F)** combined again showed that additive copies enhance combinatorial cargo partitioning. For all plots, n = 100 cells and the median partition ratios are denoted.

Next, we added in a series of equal repeats of EP tags to all three fluorescent proteins to encourage multicomponent assembly (Figure 7A, bottom). Confocal imaging of designs again confirmed that G_0_R_0_B_0_ showed no signs of visual foci (condensation or aggregation) (Figure 7D). Calculating the partition ratio likewise gave values of near 1.0 (Figure 7E). We also calculated a summed partition ratio (Figure 7F) which is the sum of all background subtracted foci signals divided by the sum of the background subtracted cellular background. For the G_0_R_0_B_0_ design, this value again came to 1.0 indicating no partitioning without EPs. Noticeable partitioning began to occur with the G_1_R_1_B_1_ designs (Figure 7D) which correlated with increased partitioning of the individual (Figure 7E) and total (Figure 7F) components. The partitioning becomes even more pronounced in cells expressing the G_2_R_2_B_2_ designs, with cell exhibiting a highly overlapping zebra-strip patterning (Figure 7D). The individual partitioning of components within the G_2_R_2_B_2_ expressing cells was not equal (Figure 7E) but followed the same trends as the G_1_R_1_B_1_, suggesting that the characteristics of the individual proteins themselves can influence partitioning, as we found that mScarlet-I3 cargo shows a moderately increased EP-dependent partitioning ability compared to mNeonGreen (Supplemental Figure 5). However, the total partitioning continued to increase (Figure 7F) and suggests that the total sum, or concentration, of EP domains drives overall partitioning.

## Discussion

### Study summary

This study focused on defining the ability of EP domains to drive self-condensation of fused cargo proteins. We found that EPs resulted in cargo partitioning both *in vivo* and *in vitro* using predictive computational simulations and wet lab experiments. This self-assembly depended on hydrophobic packing and salt bridge interactions. These interactions sum to a preference and specificity for trimeric associations. The sequence determinants are conserved among metabolosome EPs but not CcmN from carboxysomes, resulting in CcmN’s inability to promote self-condensation. Further, cargo partitioning was entirely modular and solely dependent on the inclusion of the EP domain.

### Critical residues in EP self-assembly

Our data elucidates design principles for how EPs self-assemble. Prior work had found that the hydrophobic core is critical to encapsulation within microcompartment shells,^32^ and we confirm that same propensity applies for EP self-assembly too. In addition, we identify novel salt bridges that also stabilize EP self-assembly. These salt bridges can form in a multitude of arrangements - our MD simulations suggest a preference for trimeric associations for the PduP EP. Salt bridge formation can form between various residues, including E7 and R11, in several different conformations. Our *in vitro* data found that, among the several PduP EP mutants we screened, I10 was the most critical residue in enabling condensation. Mutation of that central isoleucine nearly removed all condensation ability while disruptions to salt bridge related residues only moderately decreased condensation propensity and demonstrate that salt bridge formation alone is not sufficient to drive EP associations. As such, we propose that EP self-assembly is initially enhanced by non-specific hydrophobic packing which then enables accessory salt bridge formation for further stability and specificity. These findings refines conventional notions where EPs are regarded as any short amphipathic α-helices separated from cargo by a flexible and poorly conserved linker within a microcompartment context.^29,30^ Instead, we identify sequence elements that accommodate specific higher order networking.

### Packaging peptides in function but not in form

We tested the self-assembly of several different commonly cited encapsulation peptides from PduD, PduE, PduP, EutC, and CcmN.^28,30,46,52-54^ We were motived by the notion that β-carboxysome EPs (encoded on CcmN) act strictly as intermediaries between the procarboxysome and the shell while metabolosome EPs (encoded on PduDEP, EutC) reportedly act as a single module to aggregate and encapsulate cargo. Further, CcmN is present in very low copy numbers in β-carboxysomes^55^ while metabolosome EPs dominate interior interactions between themselves and the shell.^56^ We noted that the sequence motifs in metabolosome EPs found to be important were missing from CcmN. We provide experimental evidence that the CcmN EP does not result in clear condensation under the same conditions as metabolosome EPs, which is congruent with its physiological role as a low-occupancy tether.^26,55^ Our imaging also demonstrates metabolosome EPs can broadly trigger biomolecular condensation of fused cargo. This data makes us believe that, while the CcmN EP may be able to guide cargo encapsulation into synthetic β-carboxysome shells,^57^ it cannot be used to guide robust cargo packaging alone since it lacks the ability to self-assemble. Using metabolosome EPs, in contrast, would likely be a more robust packaging strategy for building synthetic microcompartments or shell-free enzyme partitions. “Encapsulation peptides” are still an apt name for both metabolosome and β-carboxysome EPs given their ability to package cargo within BMCs. However, a sub-categorization of “assembly peptides” may be more appropriate for metabolosome EPs which have a second role in self-assembly and could better reflect their functional diversity. In any case, an updated and more rigorous bioinformatic investigation into EP domains to better grasp their functional landscape is warranted.

Subtle differences also exist within metabolosome EPs. We observed that all our selected EPs could trigger biomolecular condensation, but the success varied (Figure 4B). The EP from PduD is found to be the most robust, followed by PduP, EutC, and PduE. This agrees with recent observations that PduCDE, led by PduD EP, triggers hierarchical assembly of accessory cargo components in Pdu BMCs.^34^ Importantly, comparing the EPs from PduD and PduP shows that the relative positioning of oppositely charged residues can be swapped as long as they maintain a sequence distance of one helical repeat. Additionally, the weak ability of the PduE EP to trigger self-condensation mirrors its prior described inability to package cargo within microcompartments and that these two functions may be related.^46^ While our data presented cannot explicitly explain the differences between different EPs, we speculate that it may relate to helical stability by flanking residues. For instance, circular dichroism of synthetic produced PduD and PduP EPs has shown that the PduD is more helical in conformation.^37^ A more stable helix could increase the entropic contribution during binding as we have observed in simulations on the PduP EP. As such, rationally designing stable helices, in conjunction with the critical sequence elements we identify, may provide a strategy for producing novel EPs with tunable properties.^32^

### Speculation on the physiological role of EP-driven condensation

One surprising finding was that the organization of EP valency (intrinsic versus emergent) drastically changes cargo partitioning and can mediate their material state *in vivo*. Serial additions of EPs onto monomeric mNeonGreen (intrinsic valency) resulted in fluid foci with moderate partitioning abilities while a single EP on tetrameric Crimson (emergent valency) turbocharged partitioning and significantly lowered liquidity, demonstrated by dramatically slower photobleaching recovery *in vivo*. Our EP-Crimson design exemplifies emergent valency that is more reflective of all characterized EP-fused enzymes. These findings suggest that BMC cargoes, which demonstrate emergent valency, can self-nucleate at a low saturation concentration and sequester nearly all cargo components during biogenesis. In native instances, the size of these assemblies may be determined by the co-expression of distinct shell proteins playing a kinetic race to truncate the growing cargo globule.^58,59^ In the propanediol microcompartment specifically, the shell factors PduABK have all been strongly linked to cargo packaging with interactions supported along their interior-presenting convex surfaces.^31,37,56,60-62^ Cargo valency and domain topology may be key considerations when packaging novel cargoes to ensure robust network formation in addition to shell targeting and packaging. While any one single EP does not adhere to the canonical stickers and spacers architecture, higher order cargo oligomers (dimers, tetramers) act as an emergent topological mimic thereof. In addition, we cannot disregard that an *in vivo* environment would modulate the self-assembly properties because due to a higher viscosity of the intracellular environment.

Our experimentation shows that salt bridge interactions, while not the predominant factor, help guide EP condensation. Knocking out the salt bridge makes cargo condensation both more sensitive to NaCl and lowers overall condensation propensity (Figure 4C, 4D). These results suggest that EP-driven cargo nucleation events may be environmentally responsive to cellular conditions, including ions. This would not be without precedent, as biomolecular condensates have been found to be regulated by metabolites, ions, and environmental conditions.^22,63,64^ Additionally, condensates themselves are recently being appreciated as also selecting for metabolites and small molecules.^65^ Taken together, these notions suggest that prometabolosomes and EP-driven condensates could partition, and be influenced by, specific cofactors and metabolites during biogenesis.^66-68^ Environmental responsiveness to redox status has been observed for CcmM-driven condensation^12^ in β-carboxysome biogenesis but further investigations for metabolosomes are warranted.

### Applications and concluding remarks

Metabolosomes can package an array of enzymatic functions within a wide array of microbes. Nature has supported this functional diversity by evolving a function agnostic, one-size-fits-all assembly approach. At its core lies the structural and functional features of the EP domain, which can be leveraged as a molecular tool to assemble disparate cargos *in vitro* and *in vivo* in a controlled, predictable manner. *In vivo*, these complexes may be used to scaffold target metabolic pathways for improved catalytic efficiencies (for example, substrate channeling). The applications for *in vitro* EP-driven phase separation are more diverse, for example: encapsulation of therapeutic molecules, stabilization of cell-free biomanufacturing systems, self-healing materials, among others. This work will help spur new hypotheses on modes of assembly and rational design efforts using EP biotechnology for novel applications.

## Acknowledgements

The authors would like to thank Dr. Eric Young for his thoughtful commentary and feedback on this manuscript. This research used resources produced by the Los Alamos National Laboratory (LANL) Institutional Computing Program, which is supported by the U.S. Department of Energy National Nuclear Security Administration under Contract No. 89233218CNA000001. The authors gratefully acknowledge the Laboratory Directed Research and Development (LDRD) program of LANL under project number 2024001DR.

## Author Contributions

**<u>D.S.T.:</u>** conceptualization, investigation, methodology, visualization, writing – original draft, writing – review and editing. **<u>C.A.L.:</u>** investigation, methodology, visualization, writing – original draft, writing - review and editing. **<u>E. R.:</u>** investigation, methodology, writing – original draft, writing – review and editing. **<u>B.L.M.:</u>** conceptualization, funding acquisition, project administration, supervision, writing – original draft, writing – review and editing. **<u>C.R.G.E.:</u>** conceptualization, project management, supervision, writing – original draft, writing – review and editing.

## STAR Methods

### Molecular Biology

A complete list of primers, synthetic gene fragments, and plasmid names used in this study can be found in the Supplemental Materials. Briefly, all cloning used the NEBuilder® HiFi DNA Assembly Cloning Kit (#E5520S), Q5® High-Fidelity 2X Master Mix (#M0492S) and/or KLD Enzyme Mix (#M0554S) offerings from New England Biolabs. Gibson assemblies were carried out using 0.02 pmol of backbone with a 3-fold molar excess of insert. All gene fragments were ordered from Twist Biosciences and codon optimized for *E. coli* and assembled into a pre-linearized pET11a backbone. Reactions were completed at 50°C for 20 minutes and then immediately frozen at -20°C until transformed into NEB5alpha chemically competent *E. coli* (#C2988J). Clone screening was performed on miniprepped plasmid DNA with whole-plasmid sequencing offered by Plasmidsaurus. PCR reactions were performed at 50 µL scale using a 15 s/kb extension time and appropriate primer annealing parameters. PCR reactions were treated with 1 µL of DpnI (#R0176S) to remove template DNA for 30 minutes at 37°C followed by cleanup and concentration using the Zymo DNA Clean and Concentrator kit and quantified with a Qubit dsDNA BR kit. For PCR mutagenesis, 1 µL of successful PCR was mixed with KLD Enzyme Mix according to manufacturer’s protocols prior to transformation.

### Protein Expression and Purification

Appropriate plasmids were transformed fresh into T7 Express (#C2566H) competent *E. coli*. Single colonies were used to inoculate 4 mL of 2xYT broth and carbenicillin antibiotic and grown overnight at 37°C, 250 RPM. Overnight cultures were then used to inoculate 250 – 500 mL of 2xYT supplemented with antibiotic in baffled flasks and grown to an optical density at 600 nm of ∼0.6-0.8 at 37°C, 250 RPM. At that density, cells were briefly cooled to room temperature prior to the addition of IPTG to 100 µM and continued incubation at 20°C and 250 RPM for 16-18 hours. Cells were harvested the next day via centrifugation (5000 RCF, 10 minutes) and frozen at -20°C for at least 30 minutes. Thawed pellets were then resuspended at 5 mL/g wet cell pellet in NP40 High Salt Lysis Buffer (RPI, #N32000) supplemented with 0.5 mM PMSF, 0.5 mg/mL egg white lysozyme, 5 mM MgCl_2_ and 12.5 U/mL recombinant benzonase (Syd Labs, #BP4200). The suspensions were lysed at 25°C for 30 minutes with constant agitation. Insoluble material was then removed by centrifugation (12000 RCF, 15 minutes, 4°C). The supernatants were then decanted into new tubes and stored on ice while the pellets were resuspended in an additional 10 mL of lysis buffer (as above) and incubated for an additional 20 minutes. The insoluble matter was removed as before, and the supernatants were combined. The supernatants were then incubated at room temperature with 1 mL of PureCube 100 Ni-INDIGO agarose beads equilibrated into Buffer A (20 mM Tris-HCl, 150 mM NaCl pH 8.0) on an end-over-end rocker for 30-60 minutes at room temperature. Proteins were then purified by gravity flow with 15 mL of washing (Buffer A + 25 mM imidazole) and 5-8 mL of elution (Buffer A + 500 mM imidazole). Proteins were then dialyzed into Buffer A such that remaining imidazole was <0.1 mM. In some instances, protein was dialyzed into 20 mM Tris-HCl pH 8.0 without additional salt. Proteins were then concentrated and quantified by A_280_ before normalization and flash freezing into liquid nitrogen to store for future use. SDS-PAGE profiles of purified proteins can be found in Supplemental Figure 6.

### Turbidity Assays

Turbidity assays were performed in 96-well plate format in quadruplicate and setup with the help of an Integra Assist Plus liquid handler in either standard (200 µL) or half-area (100 µL) plates. When assaying protein concentration effects on turbidity, stock sample was serially diluted into Assay Buffer (20 mM Tris-HCl, 150 mM NaCl pH 8.0) with appropriate mixing prior to adding an equal volume of Assay Buffer supplemented with 40% (w/v) PEG2k to reach a final value of 20%. When assaying the effects of crowding agent, an equal volume of PEG2k at a 2x final value in Assay Buffer was added to diluted protein solutions. Lastly, the effects of salt were assayed by first serially diluting a stock solution of 20 mM Tris-HCl, 5 M NaCl pH 8.0 with 20 mM Tris-HCl pH 8.0 before addition of appropriate protein followed and finally by addition of 40% PEG2k in Assay Buffer. In all cases, turbidity was measured at 600 nm in a Biotek Synergy plate reader.

### Fluorescence Resonance Energy Transfer

Condensation reactions were prepared at room temperature in triplicate 100 µL volumes in a 96-well half-area plate. Briefly, reactions were performed in 20 mM Tris-HCl, 150 mM NaCl, 20% (w/v) PEG2k with 20 µM mNeonGreen (donor) and 5 µM mScarlet-I3 (acceptor). Reactions were mixed via pipetting and left to equilibrate for 5 minutes before reading turbidity and FRET in a Biotek Synergy plate reader. A control sample consisting of 20 µM mNeonGreen alone was used to subtract non-specific FRET signal.

### Laser Scanning Confocal Microscopy

For *in vivo* imaging, protein expression was carried out using the same conditions as when expressing protein for purification. Samples were harvested with centrifugation (3000 RCF, 2 minutes) and resuspended in phosphate buffered saline (PBS) pH 7.4. Samples were then spotted into a prepared 1.5% agarose pad in PBS, spread and dried for 5 minutes prior to imaging on an Olympus FV3000. For *in vitro* imaging of droplets, 5 µL of each sample was sandwiched between two coverslips and imaged directly. The 405 nm laser line was used to excited mTagBFP2 and 488 nm laser line for mNeonGreen, 561 nm laser line for mScarlet-I3 and. Images were colored in ImageJ and are presented without further modification beyond cropping. ImageJ was also used for quantification of partition ratios by measuring the mean gray value of background subtracted foci and non-foci regions within cells. In all cases, n = 100 for partition ratio calculations. Photobleaching experiments, both *in vivo* and *in vitro*, were completed by imaging 5-10 seconds of pre-bleach sample at 0.5 s intervals prior to spot irradiation with the 488 nm laser for 0.5 seconds at 50% output. Images were then collected at 0.5 s intervals for 2 minutes. Bleaching quantification was performed in ImageJ.

### Computational Simulations

PduP EP models were based on the observed solution structure^31^ and modelled using Modeller,^69^ imposing a helical structure conformation. The helical structure was then equilibrated and simulated in a periodic box using the charmm36^70^ force field and the Gromacs^71^ molecular dynamics engine. Equilibrated peptides were later represented using the Martini coarse grained force field in its version 2.2. This step was used to explore the dimerization process of two independent helices and to extract equilibrated configurations which were backmapped into fully atomic structures using our developed pipeline.^72^ The process was repeated to simulate trimers, tetramers and pentamers.

All atomistic simulations followed the same protocols as provided here. Each backmapped peptide system was centered in a cubic box, hydrated and neutralized with 150 mM KCL. The size of the box was chosen to create at least 1.5 nm of padding on each side along the largest atom-atom distance of the peptide. A constant temperature of 310 K was maintained using velocity Langevin dynamics^73^ with a relaxation time of 1 ps. A constant isotropic pressure of 1 bar was maintained using the C-rescale approach with a relaxation time of 1 ps and compressibility of 4.5105 bar. Covalent bond lengths involving hydrogens were constrained using the SHAKE algorithm^74^ with a tolerance of 10-6 nm. Water molecules were rigidified with SETTLE.^75^ Lennard-Jones interactions were evaluated using a cut-off where forces smoothly decay to zero between 1.0 and 1.2 nm. Coulomb interactions were calculated using the particle-mesh Ewald (PME) approach. In total, 4 simulations were performed for each system with an aggregated time of 4 µs.

Coarse grained simulations were run as previously described.^76^ Simulations were carried out at 310 K using isotropic pressure coupling at 1 bar with a time step of 20 fs. Non-bonded Lennard-Jones potentials used a cutoff radius of 1.1 nm. The reaction field method was used for electrostatic calculations.^77^ The cutoff distance for dielectric constant was set to 1.1 nm. Within the cutoff, the dielectric constant was 15, and beyond the cutoff it was infinite. The velocity Verlet algorithm^78^ was used for integrating Newtonian equations. Temperature was controlled by a Langevin dynamics thermostat with a coupling time constant of 5 ps. Box pressure was controlled using a C-rescale method. For each CG MD production run, frames were saved every 100,000 time steps (every 2 ns.) Configurations were categorized using the gmx cluster tool using the gromos approach.

