## Supplemental Information for "A blueprint for biomolecular condensation driven by bacterial microcompartment encapsulation peptides"

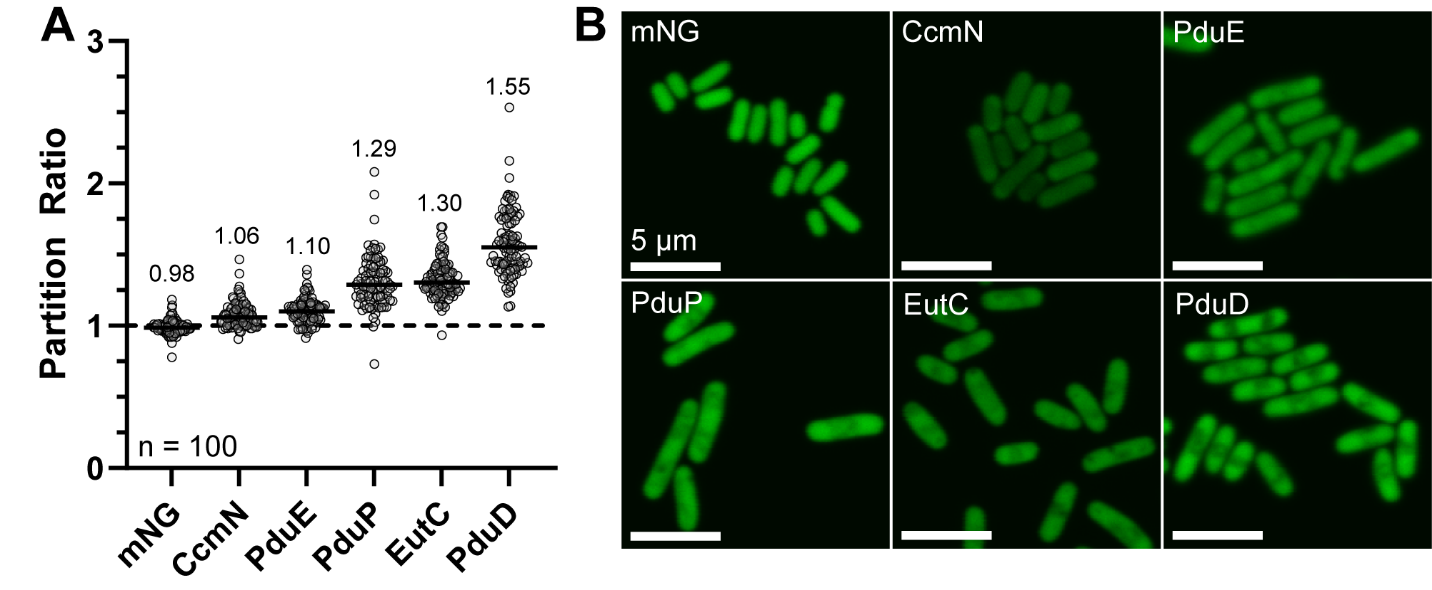

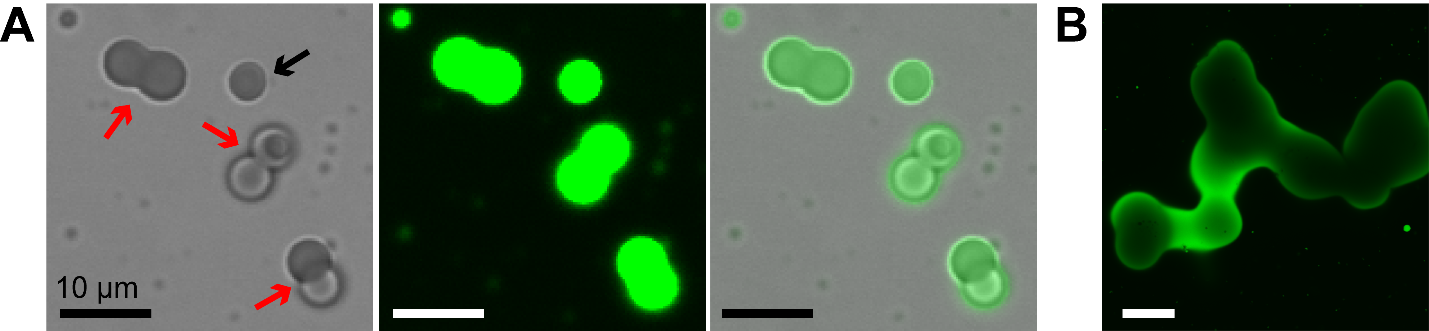
**Supplemental Figure 1: Arrested droplet fusion events. (A)** Droplets were commonly observed in mid-fusion events (red arrows) along with fully septated droplets (black arrow). **(B)** Order of mixing is critical. Adding protein last from a stock solution can lead to large aggregates that appear to be fusions of multiple droplets that could not kinetically resolve.

**Supplemental Figure 2: Morphological and functional differences for different encapsulation peptides. (A)** The partition ratio for apparent foci were calculated in cells overexpressing EP-mNeonGreen fusions. These correlate with turbidity data shown in Figure 4 in the main text. **(B)** Cells visually show variances in partitioning, with PduD being the strongest.

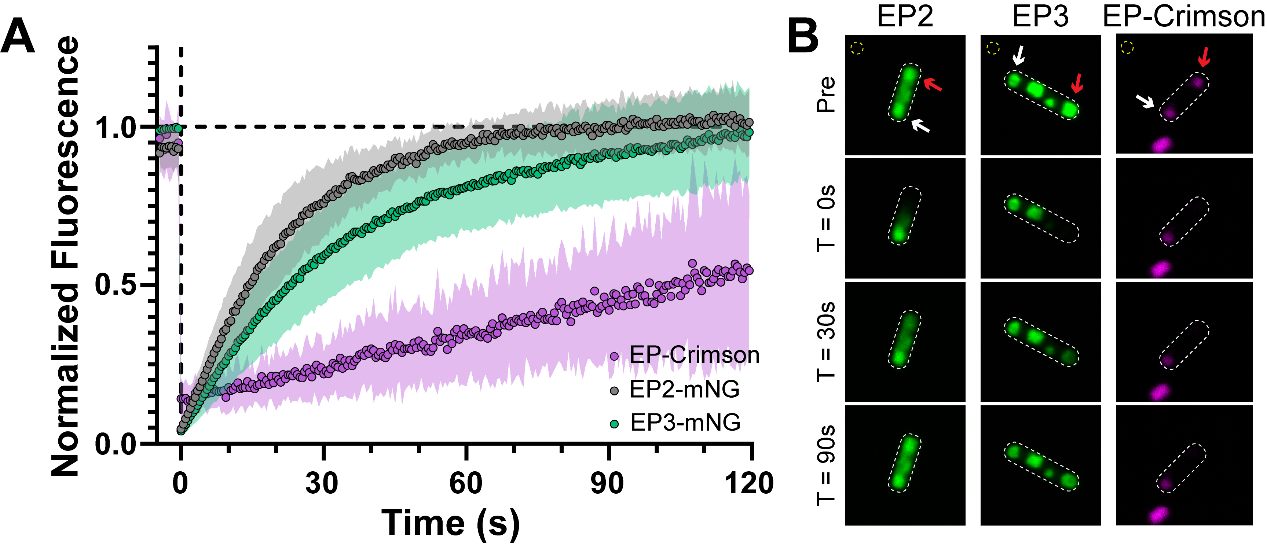

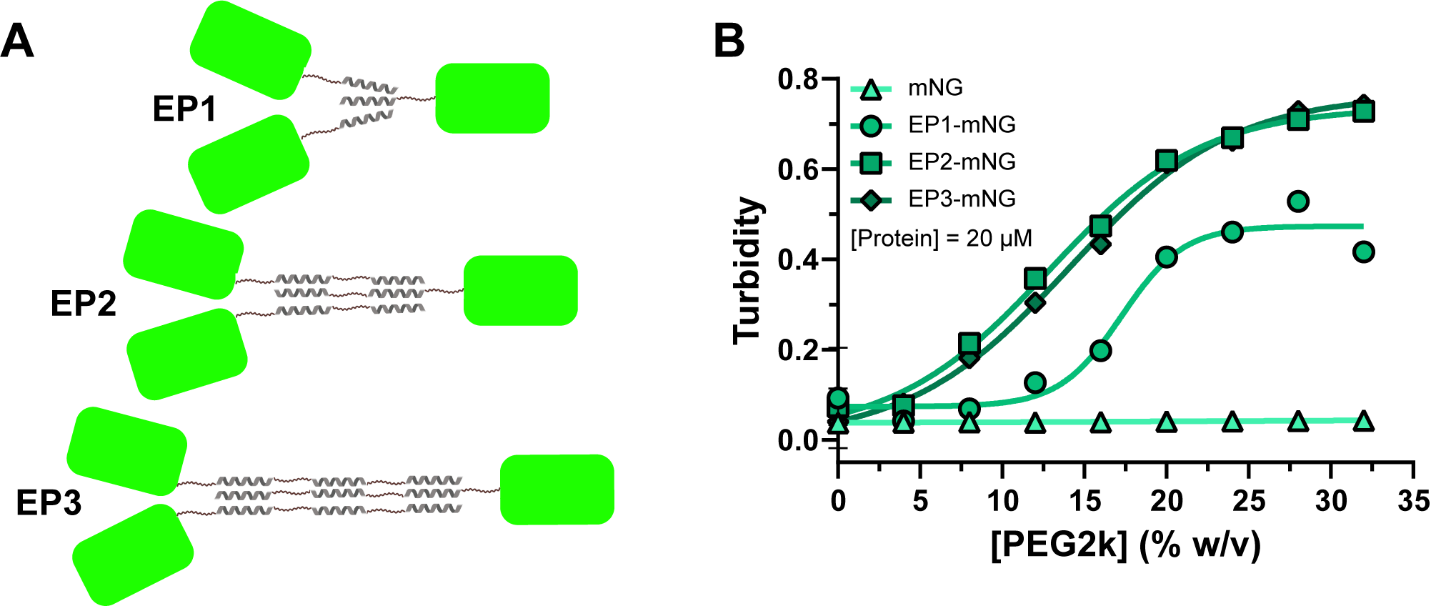
**Supplemental Figure 3: Serial additions of EPs lead to diminishing returns for enhancing condensation. (A)** Different potential configurations for binding associations of our serial EP designs, assuming a trimeric association preference. **(B)** As when titrating overall protein concentration (Figure 5A in the main text), the EP2 and EP3 designs require less PEG2k to trigger robust biomolecular condensation compared to the EP1 designs. However, the EP3 does not outperform the EP2 design, signifying self-quenching of additional networking.

**Supplemental Figure 4: Differences in material state for encapsulation peptide fused cargo depending on valency topology.** (A) Remake of Figure 6D in the main text with recovery data for the EP-Crimson (emergently valent) design overlays, showing a linear recovery profile compared to the mNeonGreen designs (intrinsically valent). (B) This recovery is demonstrated visually by limited reappearance of the bleached foci (red arrow).

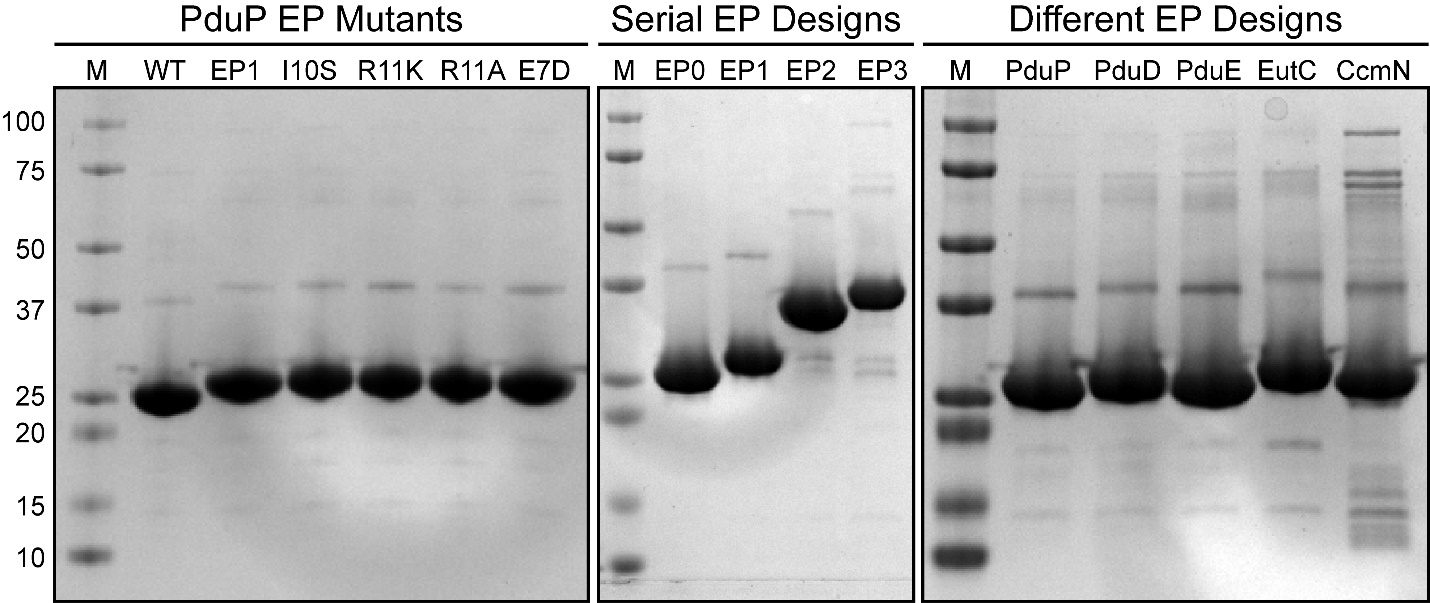

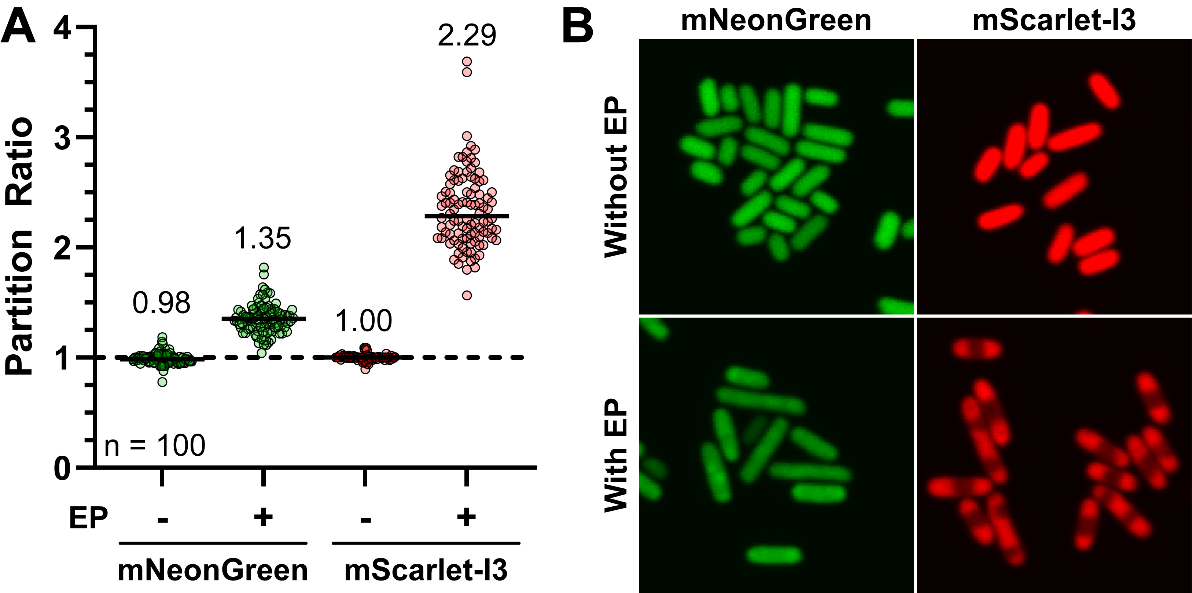
**Supplemental Figure 5:** Cargo effects on partitioning propensity. **(A)** An encapsulation peptide fused to mScarlet-I3 results in greater partitioning than the same design fused to mNeonGreen. **(B)** Confocal imaging of cells overexpressing cargo fusions show that the greater partitioning ability of mScarlet-I3 *in vivo*.

**Supplemental Figure 6: SDS-PAGE profiles of all proteins purifying and used in this study.**

**Supplemental Table 1: List of plasmids used in this study.** Acronyms used: G (mNeonGreen with C-terminal 6xHis tag), R (mScarlet-I3), and B (mTagBFP2). Subscripts refer to the number of PduP EP domains fused to the N-terminus.

| **Name** | **Source** | **Genes Encoded** | **Purpose** |
| --- | --- | --- | --- |
| pET11a | Novagen (#69436) | None | Vector backbone |
| pDTV001 | This study | G_0_ | Expression/purification |
| pDTV002 | This study | R_0_ | Expression/purification |
| pDTV004 | This study | G_1_ | Expression/purification |
| pDTV005 | This study | R_1_ | Expression/purification |
| pDTV007 | This study | G_2_ | Expression/purification |
| pDTV008 | This study | G_3_ | Expression/purification |
| pDTV009 | This study | G_0_R_0_B_0_ | Expression/purification |
| pDTV010 | This study | G_1_R_0_B_0_ | Expression/purification |
| pDTV016 | This study | G_2_R_0_B_0_ | Expression/purification |
| pDTV019 | This study | G_1_R_1_B_1_ | Expression/purification |
| pDTV026 | This study | G_3_R_0_B_0_ | Expression/purification |
| pDTV056 | This study | G_2_R_2_B_0_ | Expression/purification |
| pDTV081 | This study | R_0_-6xHis | Expression/purification |
| pDTV082 | This study | R_1_-6xHis | Expression/purification |
| pDT070 | This study | E2-Crimson-6xHis | Expression/purification |
| pDT071 | This study | EP-E2-Crimson-6xHis | Expression/purification |
| pDT101 | This study | I10S G_1_ | Expression/purification |
| pDT102 | This study | R11K G_1_ | Expression/purification |
| pDT103 | This study | R11A G_1_ | Expression/purification |
| pDT104 | This study | E7D G_1_ | Expression/purification |
| pDT107 | This study | PduD EP-mNeonGreen | Expression/purification |
| pDT108 | This study | PduE EP-mNeonGreen | Expression/purification |
| pDT109 | This study | EutC EP-mNeonGreen | Expression/purification |
| pDT110 | This study | CcmN EP-mNeonGreen | Expression/purification |

**Supplemental Table 2: List of primers used in this study.** All primers were ordered from IDT with standard desalting.

| **Name** | **Sequence (5’->3’)** | **Purpose** |
| --- | --- | --- |
| oDT122 | CGAATTCGCTAGCCCAAAAAGTTAAACAAAATTATTTCTAGAGGGGAATTG | Linearize pET11a |
| oDT123 | ATGAGAGAAGATTTTCAGCCTGACTGAGCAATAACTAGCATAACCC | Linearize pET11a |
| oDT168 | cagggtttcCAGttcGCTg | Make I10S mutation |
| oDT169 | AGCcgcaccattctgagcGAAC | Make I10S mutation |
| oDT170 | aatcagggtttcCAGttcGC | Make R11K mutation |
| oDT171 | AAAaccattctgagcGAACAGc | Make R11K mutation |
| oDT172 | GCGaccattctgagcGAACAGc | Make R11A mutation |
| oDT173 | CAGttcGCTggtGTTcatttag | Make E7D mutation |
| oDT174 | GATaccctgattcgcaccattc | Make E7D mutation |

**Supplemental Table 3: List of gene fragments used in this study.** All gene fragments were ordered from Twist Biosciences with adaptors off. Underlined sequences are the ribosomal binding site (BBa_B0034), highlighted are the gene reading frames, and bolded are the Gibson assembly overlap regions. Blue bolded overlaps will overlap directly with pET11a when linearized with oDT122 and oDT123, while red and green bolded overlaps are compatible. Fragments were designed for either single (S) assembly or multicomponent (M) assembly.

| **Name** | **Gene Encoded** | **Sequence (5’->3’)** |
| --- | --- | --- |
| gDT022 | G_0_ (S) | **TTTTTGGGCTAGCGAATTCG**tactagagaaagaggagaaatactaaATGGTGAGCAAAGGCGAAGAAGATAACATGGCGAGCCTGCCGGCGACCCATGAACTGCATATTTTTGGCAGCATTAACGGCGTGGATTTTGATATGGTGGGCCAGGGCACCGGCAACCCGAACGATGGCTATGAAGAACTGAACCTGAAAAGCACCAAAGGCGATCTGCAGTTTAGCCCGTGGATTCTGGTGCCGCATATTGGCTATGGCTTTCATCAGTATCTGCCGTATCCGGATGGCATGAGCCCGTTTCAGGCGGCGATGGTGGATGGCAGCGGCTATCAGGTGCATCGCACCATGCAGTTTGAAGATGGCGCGAGCCTGACCGTGAACTATCGCTATACCTATGAAGGCAGCCATATTAAAGGCGAAGCGCAGGTGAAAGGCACCGGCTTTCCGGCGGATGGCCCGGTGATGACCAACAGCCTGACCGCGGCGGATTGGTGCCGCAGCAAAAAAACCTATCCGAACGATAAAACCATTATTAGCACCTTTAAATGGAGCTATACCACCGGCAACGGCAAACGCTATCGCAGCACCGCGCGCACCACCTATACCTTTGCGAAACCGATGGCGGCGAACTATCTGAAAAACCAGCCGATGTATGTGTTTCGCAAGACCGAACTGAAACATAGCAAAACGGAGCTGAACTTTAAAGAATGGCAGAAAGCGTTTACCGATGTGATGGGCATGGATGAACTGTATAAAGGCGGTAGCGGCGGTAGCCACCATCACCATCACCATTAA**ATGAGAGAAGATTTTCAGCCTGA** |
| gDT023 | R_0_ (S) | **TTTTTGGGCTAGCGAATTCG**tactagagaaagaggagaaatactaaATGGATAGCACCGAAGCGGTGATTAAAGAATTTATGCGCTTTAAAGTGCATATGGAAGGCAGCATGAACGGCCATGAATTTGAAATTGAAGGCGAAGGCGAAGGCCGCCCGTATGAAGGCACCCAGACCGCGAAACTGAAAGTGACCAAAGGCGGCCCGCTGCCGTTTAGCTGGGATATTCTGAGCCCGCAGTTTATGTATGGCAGCCGCGCGTTTATTAAACATCCGGCGGATATTCCGGATTATTGGAAACAGAGCTTTCCGGAAGGCTTTAAATGGGAACGCGTGATGATTTTTGAAGATGGCGGCACCGTGAGCGTGACCCAGGATACCAGCCTGGAAGATGGCACCCTGATTTATAAAGTGAAACTGCGCGGCGGCAACTTTCCGCCGGATGGCCCGGTGATGCAGAAACGCACCATGGGCTGGGAAGCGAGCACCGAACGCCTGTATCCGGAAGATGTGGTGCTGAAAGGCGATATTAAAATGGCGCTGCGCCTGAAAGATGGCGGCCGCTATCTGGCGGATTTTAAAACCACCTATAAAGCGAAAAAACCGGTGCAGATGCCGGGCGCGTTTAACATTGATCGCAAACTGGATATTACCAGCCATAACGAAGATTATACCGTGGTGGAACAGTATGAACGCAGCGTGGCGCGCCATAGCACCGGCGGCAGCGGCGGCAGCTAA**ATGAGAGAAGATTTTCAGCCTGA** |
| gDT025 | G_1_ (S) | **TTTTTGGGCTAGCGAATTCG**tactagagaaagaggagaaatactaaatgAACaccAGCgaaCTGgaaaccctgattcgcaccattctgagcGAACAGctgaccACCccggcgCAGACCccgGTGcagccgcagggcAAAGGCattTTTcagAGCGTGAGCAAAGGCGAAGAAGATAACATGGCGAGCCTGCCGGCGACCCATGAACTGCATATTTTTGGCAGCATTAACGGCGTGGATTTTGATATGGTGGGCCAGGGCACCGGCAACCCGAACGATGGCTATGAAGAACTGAACCTGAAAAGCACCAAAGGCGATCTGCAGTTTAGCCCGTGGATTCTGGTGCCGCATATTGGCTATGGCTTTCATCAGTATCTGCCGTATCCGGATGGCATGAGCCCGTTTCAGGCGGCGATGGTGGATGGCAGCGGCTATCAGGTGCATCGCACCATGCAGTTTGAAGATGGCGCGAGCCTGACCGTGAACTATCGCTATACCTATGAAGGCAGCCATATTAAAGGCGAAGCGCAGGTGAAAGGCACCGGCTTTCCGGCGGATGGCCCGGTGATGACCAACAGCCTGACCGCGGCGGATTGGTGCCGCAGCAAAAAAACCTATCCGAACGATAAAACCATTATTAGCACCTTTAAATGGAGCTATACCACCGGCAACGGCAAACGCTATCGCAGCACCGCGCGCACCACCTATACCTTTGCGAAACCGATGGCGGCGAACTATCTGAAAAACCAGCCGATGTATGTGTTTCGCAAGACCGAACTGAAACATAGCAAAACGGAGCTGAACTTTAAAGAATGGCAGAAAGCGTTTACCGATGTGATGGGCATGGATGAACTGTATAAAGGCGGTAGCGGCGGTAGCCACCATCACCATCACCATTAA**ATGAGAGAAGATTTTCAGCCTGA** |
| gDT026 | R_1_ (S) | **TTTTTGGGCTAGCGAATTCG**tactagagaaagaggagaaatactaaatgAACaccAGCgaaCTGgaaaccctgattcgcaccattctgagcGAACAGctgaccACCccggcgCAGACCccgGTGcagccgcagggcAAAGGCattTTTcagAGCGATAGCACCGAAGCGGTGATTAAAGAATTTATGCGCTTTAAAGTGCATATGGAAGGCAGCATGAACGGCCATGAATTTGAAATTGAAGGCGAAGGCGAAGGCCGCCCGTATGAAGGCACCCAGACCGCGAAACTGAAAGTGACCAAAGGCGGCCCGCTGCCGTTTAGCTGGGATATTCTGAGCCCGCAGTTTATGTATGGCAGCCGCGCGTTTATTAAACATCCGGCGGATATTCCGGATTATTGGAAACAGAGCTTTCCGGAAGGCTTTAAATGGGAACGCGTGATGATTTTTGAAGATGGCGGCACCGTGAGCGTGACCCAGGATACCAGCCTGGAAGATGGCACCCTGATTTATAAAGTGAAACTGCGCGGCGGCAACTTTCCGCCGGATGGCCCGGTGATGCAGAAACGCACCATGGGCTGGGAAGCGAGCACCGAACGCCTGTATCCGGAAGATGTGGTGCTGAAAGGCGATATTAAAATGGCGCTGCGCCTGAAAGATGGCGGCCGCTATCTGGCGGATTTTAAAACCACCTATAAAGCGAAAAAACCGGTGCAGATGCCGGGCGCGTTTAACATTGATCGCAAACTGGATATTACCAGCCATAACGAAGATTATACCGTGGTGGAACAGTATGAACGCAGCGTGGCGCGCCATAGCACCGGCGGCAGCGGCGGCAGCTAA**ATGAGAGAAGATTTTCAGCCTGA** |
| gDT028 | G_2_ (S) | **TTTTTGGGCTAGCGAATTCG**tactagagaaagaggagaaatactaaatgAACaccAGCgaaCTGgaaaccctgattcgcaccattctgagcGAACAGctgaccACCccggcgCAGACCccgGTGcagccgcagggcAAAGGCattTTTcagAGCAACacgAGCgaaCTGgagaccctgatccgcacgattctgtcgGAACAActgaccACGccggcgCAAACCcctGTGcagcctcagggtAAAGGCattTTCcagAGCGTGAGCAAAGGCGAAGAAGATAACATGGCGAGCCTGCCGGCGACCCATGAACTGCATATTTTTGGCAGCATTAACGGCGTGGATTTTGATATGGTGGGCCAGGGCACCGGCAACCCGAACGATGGCTATGAAGAACTGAACCTGAAAAGCACCAAAGGCGATCTGCAGTTTAGCCCGTGGATTCTGGTGCCGCATATTGGCTATGGCTTTCATCAGTATCTGCCGTATCCGGATGGCATGAGCCCGTTTCAGGCGGCGATGGTGGATGGCAGCGGCTATCAGGTGCATCGCACCATGCAGTTTGAAGATGGCGCGAGCCTGACCGTGAACTATCGCTATACCTATGAAGGCAGCCATATTAAAGGCGAAGCGCAGGTGAAAGGCACCGGCTTTCCGGCGGATGGCCCGGTGATGACCAACAGCCTGACCGCGGCGGATTGGTGCCGCAGCAAAAAAACCTATCCGAACGATAAAACCATTATTAGCACCTTTAAATGGAGCTATACCACCGGCAACGGCAAACGCTATCGCAGCACCGCGCGCACCACCTATACCTTTGCGAAACCGATGGCGGCGAACTATCTGAAAAACCAGCCGATGTATGTGTTTCGCAAGACCGAACTGAAACATAGCAAAACGGAGCTGAACTTTAAAGAATGGCAGAAAGCGTTTACCGATGTGATGGGCATGGATGAACTGTATAAAGGCGGTAGCGGCGGTAGCCACCATCACCATCACCATTAA**ATGAGAGAAGATTTTCAGCCTGA** |
| gDT029 | G_3_ (S) | **TTTTTGGGCTAGCGAATTCG**tactagagaaagaggagaaatactaaatgAACaccAGCgaaCTGgaaaccctgattcgcaccattctgagcGAACAGctgaccACCccggcgCAGACCccgGTGcagccgcagggcAAAGGCattTTTcagAGCAACacgAGCgaaCTGgagaccctgatccgcacgattctgtcgGAACAActgaccACGccggcgCAAACCcctGTGcagcctcagggtAAAGGCattTTCcagAGCAATactAGCgagCTGgagactctgattcgtactattctgtcgGAGCAGctgacgACCccggcaCAGACCccgGTTcagcctcaaggcAAGGGTattTTTcagAGCGTGAGCAAAGGCGAAGAAGATAACATGGCGAGCCTGCCGGCGACCCATGAACTGCATATTTTTGGCAGCATTAACGGCGTGGATTTTGATATGGTGGGCCAGGGCACCGGCAACCCGAACGATGGCTATGAAGAACTGAACCTGAAAAGCACCAAAGGCGATCTGCAGTTTAGCCCGTGGATTCTGGTGCCGCATATTGGCTATGGCTTTCATCAGTATCTGCCGTATCCGGATGGCATGAGCCCGTTTCAGGCGGCGATGGTGGATGGCAGCGGCTATCAGGTGCATCGCACCATGCAGTTTGAAGATGGCGCGAGCCTGACCGTGAACTATCGCTATACCTATGAAGGCAGCCATATTAAAGGCGAAGCGCAGGTGAAAGGCACCGGCTTTCCGGCGGATGGCCCGGTGATGACCAACAGCCTGACCGCGGCGGATTGGTGCCGCAGCAAAAAAACCTATCCGAACGATAAAACCATTATTAGCACCTTTAAATGGAGCTATACCACCGGCAACGGCAAACGCTATCGCAGCACCGCGCGCACCACCTATACCTTTGCGAAACCGATGGCGGCGAACTATCTGAAAAACCAGCCGATGTATGTGTTTCGCAAGACCGAACTGAAACATAGCAAAACGGAGCTGAACTTTAAAGAATGGCAGAAAGCGTTTACCGATGTGATGGGCATGGATGAACTGTATAAAGGCGGTAGCGGCGGTAGCCACCATCACCATCACCATTAA**ATGAGAGAAGATTTTCAGCCTGA** |
| gDT030 | G_0_ (M) | **TTTTTGGGCTAGCGAATTCG**tactagagaaagaggagaaatactaaATGGTGAGCAAAGGCGAAGAAGATAACATGGCGAGCCTGCCGGCGACCCATGAACTGCATATTTTTGGCAGCATTAACGGCGTGGATTTTGATATGGTGGGCCAGGGCACCGGCAACCCGAACGATGGCTATGAAGAACTGAACCTGAAAAGCACCAAAGGCGATCTGCAGTTTAGCCCGTGGATTCTGGTGCCGCATATTGGCTATGGCTTTCATCAGTATCTGCCGTATCCGGATGGCATGAGCCCGTTTCAGGCGGCGATGGTGGATGGCAGCGGCTATCAGGTGCATCGCACCATGCAGTTTGAAGATGGCGCGAGCCTGACCGTGAACTATCGCTATACCTATGAAGGCAGCCATATTAAAGGCGAAGCGCAGGTGAAAGGCACCGGCTTTCCGGCGGATGGCCCGGTGATGACCAACAGCCTGACCGCGGCGGATTGGTGCCGCAGCAAAAAAACCTATCCGAACGATAAAACCATTATTAGCACCTTTAAATGGAGCTATACCACCGGCAACGGCAAACGCTATCGCAGCACCGCGCGCACCACCTATACCTTTGCGAAACCGATGGCGGCGAACTATCTGAAAAACCAGCCGATGTATGTGTTTCGCAAGACCGAACTGAAACATAGCAAAACGGAGCTGAACTTTAAAGAATGGCAGAAAGCGTTTACCGATGTGATGGGCATGGATGAACTGTATAAAGGCGGTAGCGGCGGTAGCCACCATCACCATCACCATTAA**AGCATAGCACAACGATAGCATT** |
| gDT031 | R_0_ (M) | **AGCATAGCACAACGATAGCATT**tactagagaaagaggagaaatactaaATGGATAGCACCGAAGCGGTGATTAAAGAATTTATGCGCTTTAAAGTGCATATGGAAGGCAGCATGAACGGCCATGAATTTGAAATTGAAGGCGAAGGCGAAGGCCGCCCGTATGAAGGCACCCAGACCGCGAAACTGAAAGTGACCAAAGGCGGCCCGCTGCCGTTTAGCTGGGATATTCTGAGCCCGCAGTTTATGTATGGCAGCCGCGCGTTTATTAAACATCCGGCGGATATTCCGGATTATTGGAAACAGAGCTTTCCGGAAGGCTTTAAATGGGAACGCGTGATGATTTTTGAAGATGGCGGCACCGTGAGCGTGACCCAGGATACCAGCCTGGAAGATGGCACCCTGATTTATAAAGTGAAACTGCGCGGCGGCAACTTTCCGCCGGATGGCCCGGTGATGCAGAAACGCACCATGGGCTGGGAAGCGAGCACCGAACGCCTGTATCCGGAAGATGTGGTGCTGAAAGGCGATATTAAAATGGCGCTGCGCCTGAAAGATGGCGGCCGCTATCTGGCGGATTTTAAAACCACCTATAAAGCGAAAAAACCGGTGCAGATGCCGGGCGCGTTTAACATTGATCGCAAACTGGATATTACCAGCCATAACGAAGATTATACCGTGGTGGAACAGTATGAACGCAGCGTGGCGCGCCATAGCACCGGCGGCAGCGGCGGCAGCTAA**TCGTTCAGTTTGGCTAACTCAT** |
| gDT032 | B_0_ (M) | **TCGTTCAGTTTGGCTAACTCAT**tactagagaaagaggagaaatactaaATGGTGAGCAAAGGCGAAGAACTGATTAAAGAAAACATGCATATGAAACTGTATATGGAAGGCACCGTGGATAACCATCATTTTAAATGCACCAGCGAAGGCGAAGGCAAACCGTATGAAGGCACCCAGACCATGCGCATTAAAGTGGTGGAAGGCGGCCCGCTGCCGTTTGCGTTTGATATTCTGGCGACCAGCTTTCTGTATGGCAGCAAAACCTTTATTAACCATACCCAGGGCATTCCGGATTTTTTTAAACAGAGCTTTCCGGAAGGCTTTACCTGGGAACGCGTGACCACCTATGAAGATGGCGGCGTGCTGACCGCGACCCAGGATACCAGCCTGCAGGATGGCTGCCTGATTTATAACGTGAAAATTCGCGGCGTGAACTTTACCAGCAACGGCCCGGTGATGCAGAAAAAAACCCTGGGCTGGGAAGCGTTTACCGAAACCCTGTATCCGGCGGATGGCGGCCTGGAAGGCCGCAACGATATGGCGCTGAAACTGGTGGGCGGCAGCCATCTGATTGCGAACGCGAAAACCACCTATCGCAGCAAAAAACCGGCGAAAAACCTGAAAATGCCGGGCGTGTATTATGTGGATTATCGCCTGGAACGCATTAAAGAAGCGAACAACGAAACCTATGTGGAACAGCATGAAGTGGCGGTGGCGCGCTATTGCGATCTGCCGAGCAAACTGGGCCATAAACTGAACTAA**ATGAGAGAAGATTTTCAGCCTGA** |
| gDT033 | G_1_ (M) | **TTTTTGGGCTAGCGAATTCG**tactagagaaagaggagaaatactaaatgAACaccAGCgaaCTGgaaaccctgattcgcaccattctgagcGAACAGctgaccACCccggcgCAGACCccgGTGcagccgcagggcAAAGGCattTTTcagAGCGTGAGCAAAGGCGAAGAAGATAACATGGCGAGCCTGCCGGCGACCCATGAACTGCATATTTTTGGCAGCATTAACGGCGTGGATTTTGATATGGTGGGCCAGGGCACCGGCAACCCGAACGATGGCTATGAAGAACTGAACCTGAAAAGCACCAAAGGCGATCTGCAGTTTAGCCCGTGGATTCTGGTGCCGCATATTGGCTATGGCTTTCATCAGTATCTGCCGTATCCGGATGGCATGAGCCCGTTTCAGGCGGCGATGGTGGATGGCAGCGGCTATCAGGTGCATCGCACCATGCAGTTTGAAGATGGCGCGAGCCTGACCGTGAACTATCGCTATACCTATGAAGGCAGCCATATTAAAGGCGAAGCGCAGGTGAAAGGCACCGGCTTTCCGGCGGATGGCCCGGTGATGACCAACAGCCTGACCGCGGCGGATTGGTGCCGCAGCAAAAAAACCTATCCGAACGATAAAACCATTATTAGCACCTTTAAATGGAGCTATACCACCGGCAACGGCAAACGCTATCGCAGCACCGCGCGCACCACCTATACCTTTGCGAAACCGATGGCGGCGAACTATCTGAAAAACCAGCCGATGTATGTGTTTCGCAAGACCGAACTGAAACATAGCAAAACGGAGCTGAACTTTAAAGAATGGCAGAAAGCGTTTACCGATGTGATGGGCATGGATGAACTGTATAAAGGCGGTAGCGGCGGTAGCCACCATCACCATCACCATTAAA**GCATAGCACAACGATAGCATT** |
| gDT034 | R_1_ (M) | **AGCATAGCACAACGATAGCATT**tactagagaaagaggagaaatactaaatgAACaccAGCgaaCTGgaaaccctgattcgcaccattctgagcGAACAGctgaccACCccggcgCAGACCccgGTGcagccgcagggcAAAGGCattTTTcagAGCGATAGCACCGAAGCGGTGATTAAAGAATTTATGCGCTTTAAAGTGCATATGGAAGGCAGCATGAACGGCCATGAATTTGAAATTGAAGGCGAAGGCGAAGGCCGCCCGTATGAAGGCACCCAGACCGCGAAACTGAAAGTGACCAAAGGCGGCCCGCTGCCGTTTAGCTGGGATATTCTGAGCCCGCAGTTTATGTATGGCAGCCGCGCGTTTATTAAACATCCGGCGGATATTCCGGATTATTGGAAACAGAGCTTTCCGGAAGGCTTTAAATGGGAACGCGTGATGATTTTTGAAGATGGCGGCACCGTGAGCGTGACCCAGGATACCAGCCTGGAAGATGGCACCCTGATTTATAAAGTGAAACTGCGCGGCGGCAACTTTCCGCCGGATGGCCCGGTGATGCAGAAACGCACCATGGGCTGGGAAGCGAGCACCGAACGCCTGTATCCGGAAGATGTGGTGCTGAAAGGCGATATTAAAATGGCGCTGCGCCTGAAAGATGGCGGCCGCTATCTGGCGGATTTTAAAACCACCTATAAAGCGAAAAAACCGGTGCAGATGCCGGGCGCGTTTAACATTGATCGCAAACTGGATATTACCAGCCATAACGAAGATTATACCGTGGTGGAACAGTATGAACGCAGCGTGGCGCGCCATAGCACCGGCGGCAGCGGCGGCAGCTAA**TCGTTCAGTTTGGCTAACTCAT** |
| gDT035 | B_1_ (M) | **TCGTTCAGTTTGGCTAACTCAT**tactagagaaagaggagaaatactaaatgAACaccAGCgaaCTGgaaaccctgattcgcaccattctgagcGAACAGctgaccACCccggcgCAGACCccgGTGcagccgcagggcAAAGGCattTTTcagAGCGTGAGCAAAGGCGAAGAACTGATTAAAGAAAACATGCATATGAAACTGTATATGGAAGGCACCGTGGATAACCATCATTTTAAATGCACCAGCGAAGGCGAAGGCAAACCGTATGAAGGCACCCAGACCATGCGCATTAAAGTGGTGGAAGGCGGCCCGCTGCCGTTTGCGTTTGATATTCTGGCGACCAGCTTTCTGTATGGCAGCAAAACCTTTATTAACCATACCCAGGGCATTCCGGATTTTTTTAAACAGAGCTTTCCGGAAGGCTTTACCTGGGAACGCGTGACCACCTATGAAGATGGCGGCGTGCTGACCGCGACCCAGGATACCAGCCTGCAGGATGGCTGCCTGATTTATAACGTGAAAATTCGCGGCGTGAACTTTACCAGCAACGGCCCGGTGATGCAGAAAAAAACCCTGGGCTGGGAAGCGTTTACCGAAACCCTGTATCCGGCGGATGGCGGCCTGGAAGGCCGCAACGATATGGCGCTGAAACTGGTGGGCGGCAGCCATCTGATTGCGAACGCGAAAACCACCTATCGCAGCAAAAAACCGGCGAAAAACCTGAAAATGCCGGGCGTGTATTATGTGGATTATCGCCTGGAACGCATTAAAGAAGCGAACAACGAAACCTATGTGGAACAGCATGAAGTGGCGGTGGCGCGCTATTGCGATCTGCCGAGCAAACTGGGCCATAAACTGAACTAA**ATGAGAGAAGATTTTCAGCCTGA** |
| gDT036 | G_2_ (M) | **TTTTTGGGCTAGCGAATTCG**tactagagaaagaggagaaatactaaatgAACaccAGCgaaCTGgaaaccctgattcgcaccattctgagcGAACAGctgaccACCccggcgCAGACCccgGTGcagccgcagggcAAAGGCattTTTcagAGCAACacgAGCgaaCTGgagaccctgatccgcacgattctgtcgGAACAActgaccACGccggcgCAAACCcctGTGcagcctcagggtAAAGGCattTTCcagAGCGTGAGCAAAGGCGAAGAAGATAACATGGCGAGCCTGCCGGCGACCCATGAACTGCATATTTTTGGCAGCATTAACGGCGTGGATTTTGATATGGTGGGCCAGGGCACCGGCAACCCGAACGATGGCTATGAAGAACTGAACCTGAAAAGCACCAAAGGCGATCTGCAGTTTAGCCCGTGGATTCTGGTGCCGCATATTGGCTATGGCTTTCATCAGTATCTGCCGTATCCGGATGGCATGAGCCCGTTTCAGGCGGCGATGGTGGATGGCAGCGGCTATCAGGTGCATCGCACCATGCAGTTTGAAGATGGCGCGAGCCTGACCGTGAACTATCGCTATACCTATGAAGGCAGCCATATTAAAGGCGAAGCGCAGGTGAAAGGCACCGGCTTTCCGGCGGATGGCCCGGTGATGACCAACAGCCTGACCGCGGCGGATTGGTGCCGCAGCAAAAAAACCTATCCGAACGATAAAACCATTATTAGCACCTTTAAATGGAGCTATACCACCGGCAACGGCAAACGCTATCGCAGCACCGCGCGCACCACCTATACCTTTGCGAAACCGATGGCGGCGAACTATCTGAAAAACCAGCCGATGTATGTGTTTCGCAAGACCGAACTGAAACATAGCAAAACGGAGCTGAACTTTAAAGAATGGCAGAAAGCGTTTACCGATGTGATGGGCATGGATGAACTGTATAAAGGCGGTAGCGGCGGTAGCCACCATCACCATCACCATTAA**AGCATAGCACAACGATAGCATT** |
| gDT037 | R_2_ (M) | **AGCATAGCACAACGATAGCATT**tactagagaaagaggagaaatactaaatgAACaccAGCgaaCTGgaaaccctgattcgcaccattctgagcGAACAGctgaccACCccggcgCAGACCccgGTGcagccgcagggcAAAGGCattTTTcagAGCAACacgAGCgaaCTGgagaccctgatccgcacgattctgtcgGAACAActgaccACGccggcgCAAACCcctGTGcagcctcagggtAAAGGCattTTCcagAGCGATAGCACCGAAGCGGTGATTAAAGAATTTATGCGCTTTAAAGTGCATATGGAAGGCAGCATGAACGGCCATGAATTTGAAATTGAAGGCGAAGGCGAAGGCCGCCCGTATGAAGGCACCCAGACCGCGAAACTGAAAGTGACCAAAGGCGGCCCGCTGCCGTTTAGCTGGGATATTCTGAGCCCGCAGTTTATGTATGGCAGCCGCGCGTTTATTAAACATCCGGCGGATATTCCGGATTATTGGAAACAGAGCTTTCCGGAAGGCTTTAAATGGGAACGCGTGATGATTTTTGAAGATGGCGGCACCGTGAGCGTGACCCAGGATACCAGCCTGGAAGATGGCACCCTGATTTATAAAGTGAAACTGCGCGGCGGCAACTTTCCGCCGGATGGCCCGGTGATGCAGAAACGCACCATGGGCTGGGAAGCGAGCACCGAACGCCTGTATCCGGAAGATGTGGTGCTGAAAGGCGATATTAAAATGGCGCTGCGCCTGAAAGATGGCGGCCGCTATCTGGCGGATTTTAAAACCACCTATAAAGCGAAAAAACCGGTGCAGATGCCGGGCGCGTTTAACATTGATCGCAAACTGGATATTACCAGCCATAACGAAGATTATACCGTGGTGGAACAGTATGAACGCAGCGTGGCGCGCCATAGCACCGGCGGCAGCGGCGGCAGCTAA**TCGTTCAGTTTGGCTAACTCAT** |
| gDT038 | B_2_ (M) | **TCGTTCAGTTTGGCTAACTCAT**tactagagaaagaggagaaatactaaatgAACaccAGCgaaCTGgaaaccctgattcgcaccattctgagcGAACAGctgaccACCccggcgCAGACCccgGTGcagccgcagggcAAAGGCattTTTcagAGCAACacgAGCgaaCTGgagaccctgatccgcacgattctgtcgGAACAActgaccACGccggcgCAAACCcctGTGcagcctcagggtAAAGGCattTTCcagAGCGTGAGCAAAGGCGAAGAACTGATTAAAGAAAACATGCATATGAAACTGTATATGGAAGGCACCGTGGATAACCATCATTTTAAATGCACCAGCGAAGGCGAAGGCAAACCGTATGAAGGCACCCAGACCATGCGCATTAAAGTGGTGGAAGGCGGCCCGCTGCCGTTTGCGTTTGATATTCTGGCGACCAGCTTTCTGTATGGCAGCAAAACCTTTATTAACCATACCCAGGGCATTCCGGATTTTTTTAAACAGAGCTTTCCGGAAGGCTTTACCTGGGAACGCGTGACCACCTATGAAGATGGCGGCGTGCTGACCGCGACCCAGGATACCAGCCTGCAGGATGGCTGCCTGATTTATAACGTGAAAATTCGCGGCGTGAACTTTACCAGCAACGGCCCGGTGATGCAGAAAAAAACCCTGGGCTGGGAAGCGTTTACCGAAACCCTGTATCCGGCGGATGGCGGCCTGGAAGGCCGCAACGATATGGCGCTGAAACTGGTGGGCGGCAGCCATCTGATTGCGAACGCGAAAACCACCTATCGCAGCAAAAAACCGGCGAAAAACCTGAAAATGCCGGGCGTGTATTATGTGGATTATCGCCTGGAACGCATTAAAGAAGCGAACAACGAAACCTATGTGGAACAGCATGAAGTGGCGGTGGCGCGCTATTGCGATCTGCCGAGCAAACTGGGCCATAAACTGAACTAA**ATGAGAGAAGATTTTCAGCCTGA** |
| gDT039 | G_3_ (M) | **TTTTTGGGCTAGCGAATTCG**tactagagaaagaggagaaatactaaatgAACaccAGCgaaCTGgaaaccctgattcgcaccattctgagcGAACAGctgaccACCccggcgCAGACCccgGTGcagccgcagggcAAAGGCattTTTcagAGCAACacgAGCgaaCTGgagaccctgatccgcacgattctgtcgGAACAActgaccACGccggcgCAAACCcctGTGcagcctcagggtAAAGGCattTTCcagAGCAATactAGCgagCTGgagactctgattcgtactattctgtcgGAGCAGctgacgACCccggcaCAGACCccgGTTcagcctcaaggcAAGGGTattTTTcagAGCGTGAGCAAAGGCGAAGAAGATAACATGGCGAGCCTGCCGGCGACCCATGAACTGCATATTTTTGGCAGCATTAACGGCGTGGATTTTGATATGGTGGGCCAGGGCACCGGCAACCCGAACGATGGCTATGAAGAACTGAACCTGAAAAGCACCAAAGGCGATCTGCAGTTTAGCCCGTGGATTCTGGTGCCGCATATTGGCTATGGCTTTCATCAGTATCTGCCGTATCCGGATGGCATGAGCCCGTTTCAGGCGGCGATGGTGGATGGCAGCGGCTATCAGGTGCATCGCACCATGCAGTTTGAAGATGGCGCGAGCCTGACCGTGAACTATCGCTATACCTATGAAGGCAGCCATATTAAAGGCGAAGCGCAGGTGAAAGGCACCGGCTTTCCGGCGGATGGCCCGGTGATGACCAACAGCCTGACCGCGGCGGATTGGTGCCGCAGCAAAAAAACCTATCCGAACGATAAAACCATTATTAGCACCTTTAAATGGAGCTATACCACCGGCAACGGCAAACGCTATCGCAGCACCGCGCGCACCACCTATACCTTTGCGAAACCGATGGCGGCGAACTATCTGAAAAACCAGCCGATGTATGTGTTTCGCAAGACCGAACTGAAACATAGCAAAACGGAGCTGAACTTTAAAGAATGGCAGAAAGCGTTTACCGATGTGATGGGCATGGATGAACTGTATAAAGGCGGTAGCGGCGGTAGCCACCATCACCATCACCATTAA**AGCATAGCACAACGATAGCATT** |
